# The Effect of Task on Object Processing revealed by EEG decoding

**DOI:** 10.1101/2020.08.18.255018

**Authors:** Hoi Ming Ken Yip, Leo Y. T. Cheung, Vince S. H. Ngan, Yetta Kwailing Wong, Alan C.-N. Wong

## Abstract

Recent studies showed that task demand affects object representations in higher-level visual areas and beyond, but not so much in earlier areas. There are, however, limitations in those studies including the relatively weak manipulation of task due to the use of familiar real-life objects, the low temporal resolution in fMRI, and the emphasis on the amount and not the source of information carried by brain activations. In the current study, observers categorized images of artificial objects in one of two orthogonal dimensions, shape and texture, while their brain activity was recorded with electroencephalogram (EEG). Results showed that object processing along the texture dimension was affected by task demand starting from a relatively late time (320-370ms time window) after image onset. The findings are consistent with the view that task exerts an effect on the later phases of object processing.

## Introduction

Object recognition and categorization depend greatly on the goal of the observer, and yet its underlying mechanism has not been well understood. On the one hand, feedforward theories portrait object processing as a hierarchy in the ventral visual pathway, with representations increasing in receptive field size, complexity, and invariance going from the earlier areas like the primary visual cortex (V1) to later areas in the inferior temporal cortex (IT) (DiCarlo et al., 2012; Grill-Spector et al., 2001; Kriegeskorte et al., 2008). Several computational models of object representation proposed that the processing from V1 to IT is largely feedforward and not influenced by top-down factors such as task (DiCarlo et al., 2012; Riesenhuber & Poggio, 2002; Serre et al., 2005; Serre et al., 2007).

On the other hand, the role of top-down factors in visual object representations is conceivable in more recent perception models with an element of predictive coding (Clark, 2013; Friston, 2010; Hohwy, 2013, 2017; Lupyan 2015; Macpherson, 2017; O’Callaghan et al., 2017; Ransom et al., 2017; Vance & Stokes, 2017). These models share the same basic principle of prediction error minimization: higher-level areas generate predictions and send them to lower-level areas, which compare the predictions with the incoming information and send the prediction errors (difference between the two) back to the higher-level areas for adjusting the predictions. This process is repeated at both high (e.g., whole object representations) and low levels (e.g., edge orientations). Crucially, low-level predictions can be influenced by top-down factors such that experience, knowledge, and expectations in the form of predictions passed to the lower-level areas, even all the way to the primary visual cortex (Kok et al., 2012; Lupyan, 2015). Accordingly, task can affect object representations even in the earlier phases of visual processing.

Previous studies have examined the effect of task demand on object representations in the visual system. Findings from some event-related potential (ERP) studies suggest that face representations are affected by task (Goffaux et al., 2003; Itier, & Neath-Tavares, 2017; Valdés-Conroy et al., 2014). For example, Goffaux et al. (2003) presented normal, low-spatial-frequency, and high-spatial-frequency faces to participants who were asked to judge the gender or familiarity of the faces. For low-spatial-frequency faces, an enhanced amplitude in the N170 was found when participants performed the gender task than the familiar task. No task effect was found for normal or high-spatial-frequency faces.

Though revealing, there are limitations in the interpretation of the findings from the traditional, activation-based approach as in the aforementioned studies. For example, in Goffaux et al. (2003), the task effect in the N170 amplitude was interpreted as enhanced diagnosticity of low-spatial-frequency cues in the gender task than the familiarity task. Nevertheless, an enhanced amplitude for one direction does not necessarily imply information diagnosticity and can be caused by either a higher neural engagement or lower processing efficiency. Also, similar N170 amplitudes for the two tasks in the high-spatial-frequency faces does not necessarily mean a lack of task effect. It is possible that the task exerts an effect not on the amplitude of the N170 in general for a few selected channels but on the pattern of activation across channels. In addition, extraneous factors such as changes in dipole position can contribute to the direction of amplitude difference.

### MVPA Studies of Task Effects

Compared with the traditional, univariate analysis, multivariate pattern analysis (MVPA) is more suited to research questions concerning the effect of task on object representations (for a detailed introduction of the technique, see Grootswagers et al., 2017; Haynes, 2015; Schwarzkopf & Rees, 2011). A specific type of MVPA is decoding, which involves the use of machine learning algorithms to infer the stimuli or experimental conditions based on the pattern of brain activations across space (e.g., channels for EEG, voxels for fMRI) and time. Classification accuracy indicates the success of such inference and thus how differently the brain responds to two stimuli or conditions. The technique is useful for showing the amount of diagnostic information carried by the spatio-temporal patterns of activations for distinguishing between different objects. Hence, the degree to which the activation patterns between objects are distinguishable under different task contexts can reflect the effect of task demand on object representations.

In recent years, several studies applied MVPA to examine task effect on object processing. The majority of these studies used functional magnetic resonance imaging (fMRI). For example, by computing the correlations of different objects in either the same task or different tasks, Harel et al. (2014) observed that object identity information differed in the posterior fusiform gyrus (pFs) but not in the lateral occipital cortex (LO). In Bracci et al. (2017), while analyses with the representational dissimilarity matrices (RDM) suggested a task effect in the ventral but not lateral occipitotemporal cortex, the correlation between the response patterns for objects remained similar regardless of whether the task was the same or different in both regions. Kim & McCarthy (2016) reported a task effect on decoding face versus body images in the occipitotemporal fusiform region, but this finding came with a counter-intuitive observation that the face-body decoding was *less* accurate when participants were performing a face-body discrimination task compared with an irrelevant emotion discrimination task. Jackson et al. (2017) reported above-chance decoding in the lateral occipital complex (LOC) only when the task was relevant, although the decoding accuracy in this task-relevant condition was not significantly different from that in the task-irrelevant condition. Two other studies did not observe any significant task effect (Long & Kuhl, 2018; Vaziri-Pashkam & Xu, 2017).

Fewer studies use techniques that are temporally more sensitive. Using magnetoencephalography (MEG), Hebart et al. (2018) observed a task effect on object decoding performance only after 542ms, suggesting that task affects object representation only at a late time. However, in another study using EEG and time-frequency analyses, decoding performance was higher in a task-relevant than a task-irrelevant condition during a much earlier time frame (100-600ms after stimulus onset), but it was the case only for one of the two tasks included in that study (Bocincova & Johnson, 2019).

Overall, previous findings do not provide conclusive evidences concerning the effect of task on object representations as well as its time-course. Three reasons may have contributed to the mixed findings. First, most studies used real-life objects (Bracci et al., 2017; Harel et al., 2014; Hebart et al., 2018; Kim and McCarthy, 2016; Long & Kuhl, 2018; Vaziri-Pashkam & Xu, 2017). In daily life, we often perform multiple tasks on the same object simultaneously (e.g. recognizing the gender and emotion of a face when you want to know if a girl is happy). It may thus be difficult for participants in those studies to focus on just one dimension as instructed, resulting in weaker manipulation of task across conditions. Consistently, the two studies using artificial stimuli (Bocincova & Johnson, 2019; Jackson et al., 2017) both found some support for the task-dependent object representations.

Second, most studies used fMRI to measure brain activation, a technique with low temporal resolution in the order of seconds. It is thus unclear *when* task starts to influence object processing. On the one hand, task effects have been observed in the brain regions corresponding to visual object processing and representation such as the ventral temporal cortex and lateral occipital cortex (Bracci et al., 2017; Harel et al., 2014; Jackson et al., 2017; Kim & McCarthy, 2016), and can be interpreted as the penetration of task into the initial formation of object representation early in time (Pylyshyn, Z., 1999). An alternative interpretation, however, is that the effects may have been caused by the late feedback of high-level processing to visual areas to modify existing object representations. Techniques with a higher temporal resolution such as EEG and MEG would be more useful in teasing apart these different interpretations. Indeed, inconsistent findings have been observed between fMRI and EEG studies on visual processing of simple features. For example, attention and expectation have been shown to modulate responses to gratings in primary visual cortex (V1) in previous fMRI studies (e.g. Jehee et al., 2011; Kok et al., 2012), but not the amplitude of C1, the earliest ERP component that originates from V1 and peaks before 100ms, in another ERP study (Alilović et al., 2019). Therefore, it is important to use a time-sensitive method to clarify this timing of the task effect.

The third potential reason of the inconclusive results in the literature concerns what exactly about object representations is affected by task. Typically, previous studies focused on the *amount of information* carried by the pattern of brain activations, and compared the decoding accuracy between task-relevant and irrelevant conditions. Better decoding accuracy in the task-relevant condition would mean that more information is carried by the pattern of brain activations for object distinctions consistent with the observer’s current task demand. Yet a lack of difference in decoding accuracy does not necessarily mean that task does not affect object representations. A much less explored possibility is that the *source of the information* useful for object distinction may differ depending on task. That is, the exact features (in terms of channels, time points, etc.) useful for object decoding might change with task. It could therefore be the case that in some of the previous studies where a task effect on decoding accuracy was not found, task actually did influence object representations in terms of which brain activation features constitute the information useful for object differentiation. Examination of both the amount and source of information carried by the brain action pattern would provide a more complete picture of the effect of task on object representations.

### The Current Study

In the current study we asked whether and when task effects start to emerge during the time course of visual object processing to clarify whether task affects object representations during or after their formation. MVPA was applied to the EEG responses of participants performing a verification task on the objects based on one of two possible dimensions (shape and texture). With respect to the limitations of previous studies reviewed above, we introduced a stronger manipulation of task by using novel, artificial objects. This should increase the likelihood that participants attended to only one dimension at a time as instructed. Besides, the use of EEG also offered a better temporal resolution compared with techniques like fMRI.

Task effect can be manifested in many ways. One approach is to ask how task affects decoding of object categories (Bracci et al. 2017; Harel et al., 2014; Hebart et al., 2018; Vaziri-Pashkam & Xu, 2017). These studies presented objects from a variety of categories and examined the classifiers’ accuracies in differentiating these objects in different tasks, typically involved categorizing objects in terms of physical (e.g. tilt, color) and conceptual properties (e.g. animacy, movement). However, as the difference between physical and conceptual tasks were mainly on the depth of semantic analysis, the perceptual processing required may actually be similar, which is not ideal if we want to maximize the effect of task on object processing. Another approach is to design stimuli with physical features varying in orthogonal dimensions and require participants to focus on one dimension in each task (Bocincova & Johnson, 2019; Jackson et al., 2017; Kim and McCarthy, 2016; Long & Kuhl, 2018). The current study adopted the latter approach, with artificial stimuli which allowed us to design orthogonal dimensions and maximize the effect of task.

In addition, we also examined both the amount and the source of the information carried by brain activations. To achieve these, two sets of MVPA analyses were performed. One set of analyses compared the decoding performance of task-relevant classifiers with that of the task-irrelevant classifiers (Table 1) to examine whether task relevance affects the amount of information useful for object differentiation. If task affects the amount of information in object representations, a higher object decoding accuracy should be observed when the decoded dimension (e.g., texture) is task-relevant than when the dimension is task-irrelevant, i.e., the participant’s task was to judge an object along the same dimension (e.g., texture) compared with a different dimension (e.g., shape). The other set of analyses concerned whether classifiers trained on a particular task context performed better when they were tested on the same task context (‘task-relevant classifiers’) than on a different task context (‘across-task classifiers’; Table 1). This directly tested how useful the features extracted during classifier training in one task context were when applied to the decoding in a different task context. If task affects the source of information carried in object representations, the decoding model trained in one task context would generalize better to data in the same task context than to a different task context, i.e., the decoding accuracy for a dimension. e.g., texture, should be better when classifier training and testing were both conducted on data in the texture task context (‘task-relevant classifiers’; Table 1) compared with when classifier was trained with data in the texture task context and then tested with data in the shape task context (‘across-task classifiers’; Table 1).

**Table 1.**
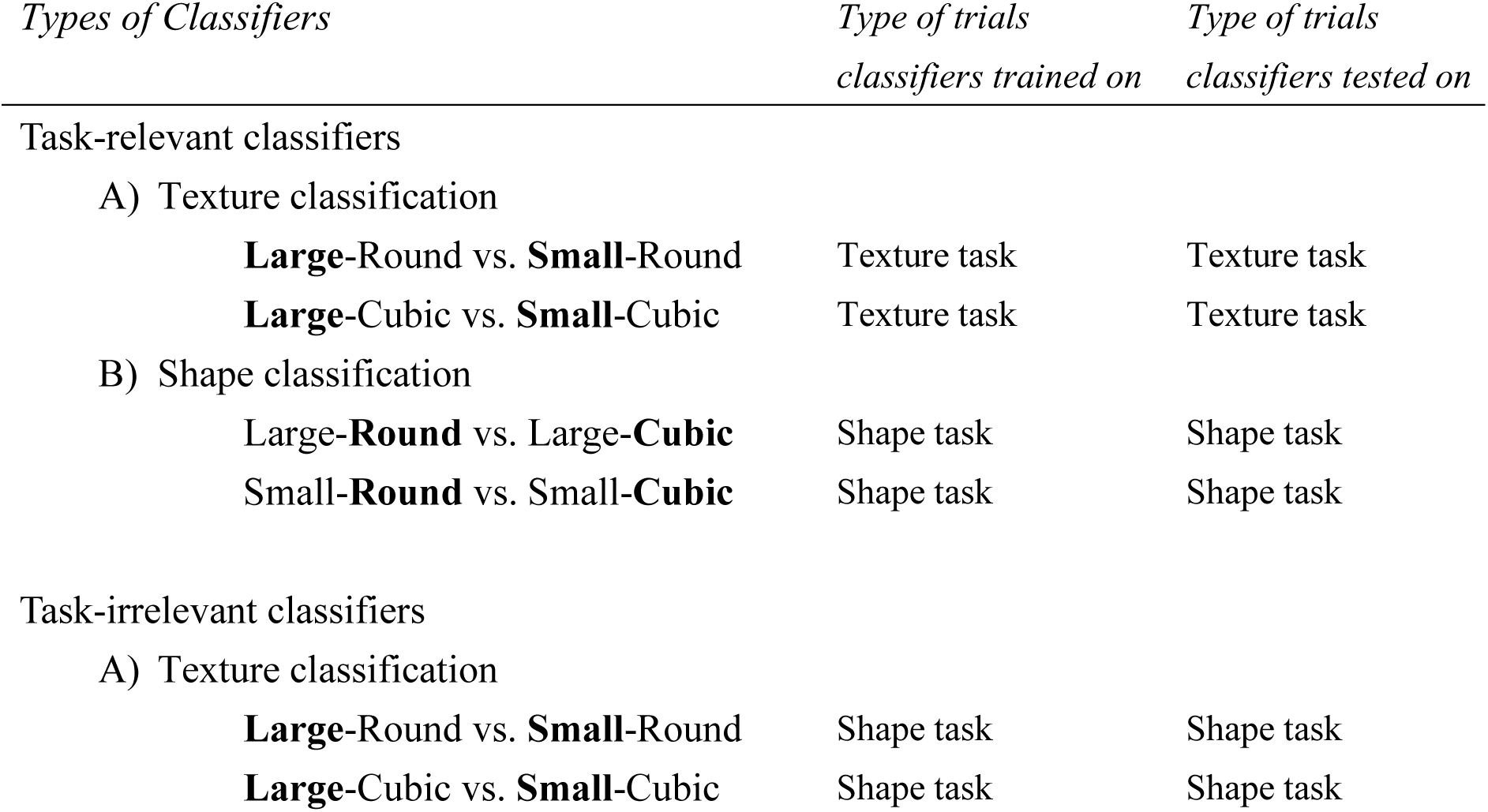
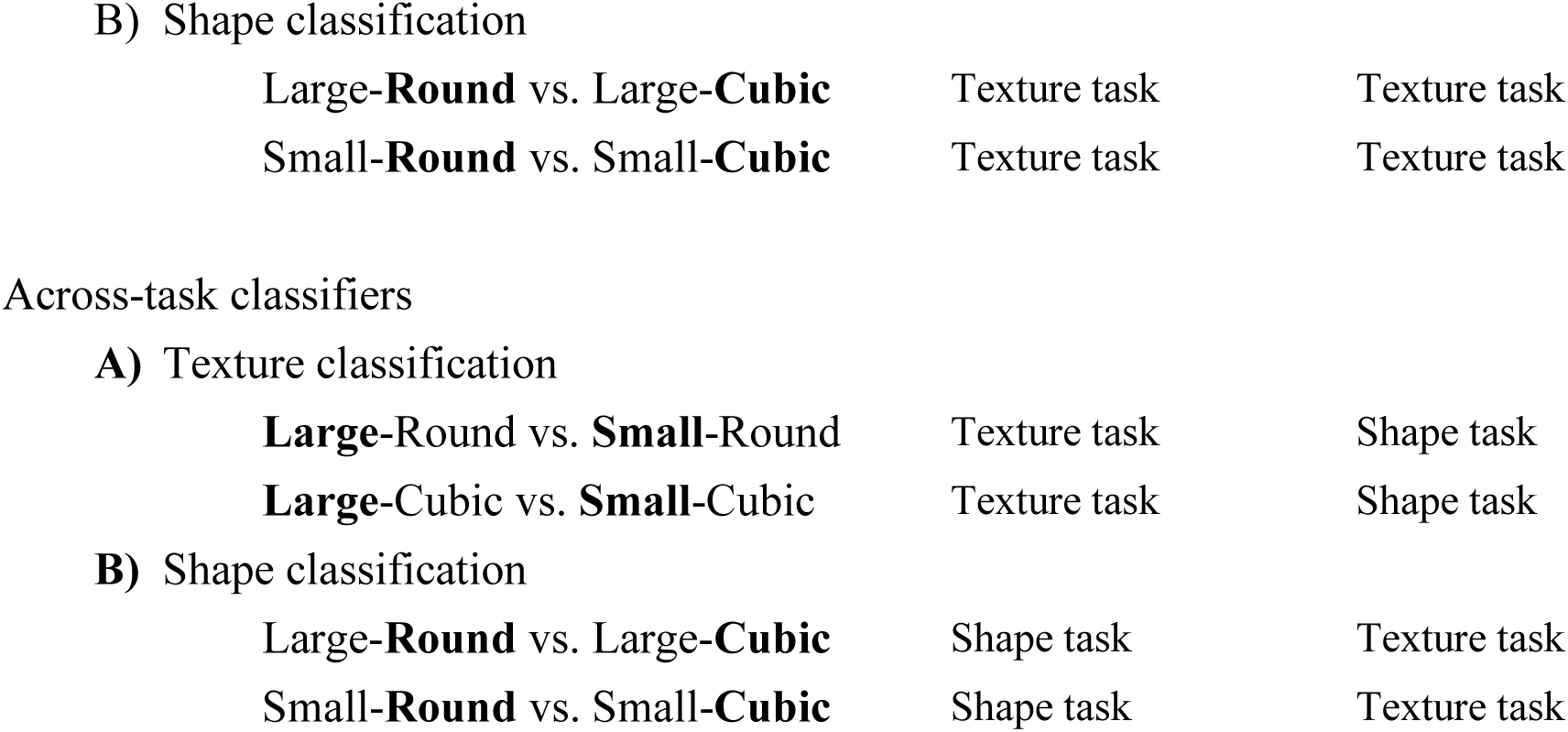
The three types of classifiers involved in the current study. Dimension values relevant to the classification were highlighted in bold.

Apart from *whether* there is any task effect on object representations, we also asked *when* task effects arise in time. If task affects the formation of object representation, task effects should be observed at or before the time range of the N170 and the N250, two important ERP components important for visual object recognition (150-330ms; Goffaux et al., 2003; Scott et al., 2006). The N170 has been associated mainly with initial coding of faces and familiar objects (Bentin et al., 1996; Scott et al., 2006; Tanaka & Curran, 2001). The N250, on the other hand, has been shown to be sensitive to the identity of individual faces and objects (Pierce et al., 2011; Schweinberger et al., 2002, Schweinberger et al., 2004; Scott et al., 2006). Source localization studies found that these components have a major source in the fusiform gyrus (Deffke et al., 2007; Herrmann et al., 2005; Schweinberger et al., 2002; Schweinberger et al., 2004; Scott et al., 2006), which correspond well with the brain regions in which task effects have been observed using fMRI (Bracci et al., 2017; Harel et al., 2014; Jackson et al., 2017; Kim & McCarthy, 2016). Alternatively, if task only modifies object representations through feedback signals from higher-level processing, then task effects should only be observed later in time than the time window of these two ERP components.

## Materials & Methods

### Participants

Forty college students with normal or corrected-to-normal vision were recruited. Four participants were excluded due to low behavioral accuracy (<90%) or excessive artifacts (>20% of total trials rejected in artifact rejection; see below). Data of the remaining 36 participants (17 males, mean age = 21.83 years, SD = 3.47) were included in all subsequent analyses. All participants gave written consent according to the guidelines of the Survey and Behavioral Research Ethics Committee of the Chinese University of Hong Kong and the Joint Chinese University of Hong Kong-New Territories East Cluster Clinical Research Ethics Committee, and received monetary compensation for their participation.

### Materials and stimuli

Artificial objects were generated and modified based on the computer-generated artificial objects (‘Greebles’ and ‘Ziggerins’) used in previous studies (Figure 1; Gauthier & Tarr, 1997; Wong et al., 2009). The objects were in an identical brown color and varied in two dimensions: shape and texture. The luminance levels were controlled across the object groups. The objects were sized such that a circle enclosing each object would span a visual angle of 4° X 4°.

**Figure 1.**
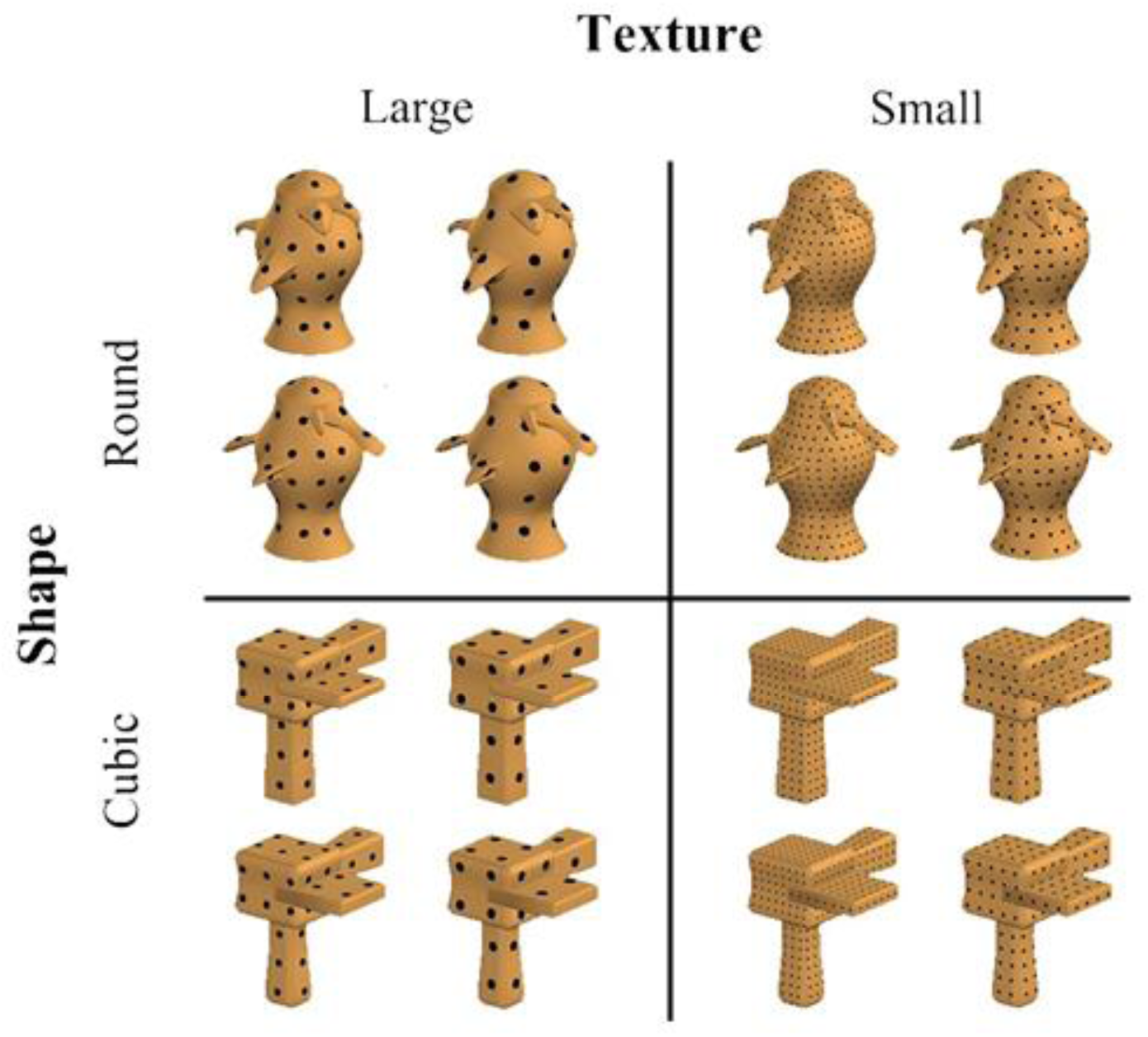
The four groups of objects varying along the shape and texture dimensions. For better visualization the objects were brighter in this figure than in the actual experiment.

A PC with Matlab (Mathworks, Inc.) and Psychophysics Toolbox extensions (Brainard, 1997; Pelli, 1997) was used to present stimuli and record behavioral responses. Stimuli were presented on a 27” LCD monitor with a refresh rate of 60Hz. The background was grey and the viewing distance was 70cm.

### Procedure

A verification task was used in this study (Figure 2). On each trial, a cue (“large”, “small”, “round” or “cubic”) was presented for 500ms, followed by a black fixation cross for 1000ms. Then an object was presented for 1000ms. Participants were asked to press “1” on the keyboard if the object matched with the cue and “2” if it did not. Trials with “large” or “small” cues required judgment about the texture (i.e., a texture task) and trials with “round” or “cubic” cues required judgment about the shape (i.e., a shape task). Participants were instructed to respond as accurately and as quickly as possible within the 1000ms when the object appeared, and the object remained present after the response. Following the presentation of the object, a white fixation cross was presented for 2000ms. Participants were instructed to blink only during this period.

**Figure 2.**
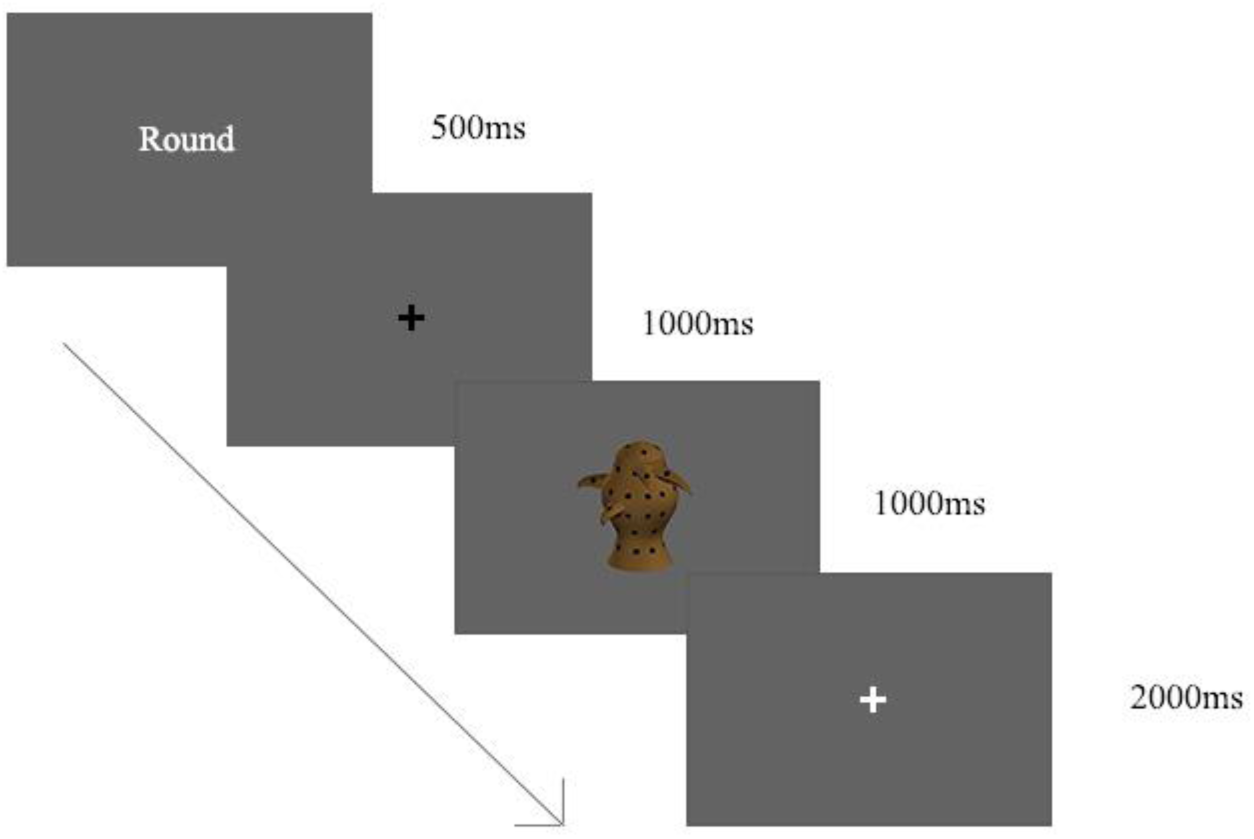
The experimental paradigm. The task was to judge whether the object matched with the cue.

There were 32 blocks and 32 trials in each block, leading to 1024 trials in total. Participants were advised by the experimenter to get at least 30 out of 32 trials correct in each block and received feedback on the number of correct trials after each block. Each mapping of cue and object groups (4 cue X 4 object groups = 16 pairs) was presented twice in a block. The selection of objects within each object group was random. The trial sequence within a block was pseudo-randomized with the following criteria: (i) The same cue did not repeat for more than 3 consecutive trials; (ii) The same object group did not repeat for more than 3 consecutive trials; Sixty-four different trial sequences were generated and 32 of them were randomly selected for each participant. There was a 12-trial short practice before the experiment. Participants were required to obtain 90% accuracy to proceed. Otherwise they would repeat the practice until they attained the required accuracy.

### EEG recording and Preprocessing

EEG data were recorded on a 128-channel eego system (ANT Neuro, Enschede, The Netherlands) with a 1000-Hz sampling rate. CPz was used as the online reference and the ground electrode was placed close to the left mastoid. Electrode impedance was kept under 20kΩ during the experiment.

Preprocessing was conducted with EEGLAB (Delorme & Makeig, 2004) and ERPLAB (Lopez-Calderon & Luck, 2014) in MATLAB (Mathworks, Natick, MA). Data were re-referenced off-line to the average reference. Band-pass filtering (0.01–30Hz, EEGLAB basic FIR filter, filter order = 66000) was applied to the continuous EEG data, which were then divided into 1000-ms epochs starting at 200ms before and ending at 800ms after the presentation of the object. Since the signals near the end of the stimulus presentation was not of interest, the last 200ms of stimuli presentation (i.e., 800-1000ms) was not used for artifact rejection to avoid rejecting epochs due to artifacts during this time window. Incorrect trials were also excluded from further analyses. Epochs with ocular artifacts were removed by visual inspection and by the moving-window peak-to-peak function in ERPLAB on VEOG, HEOG and the channels selected in the decoding analysis with a threshold of 100 μV, a window size of 200ms and a step size of 50ms. On average, 4.57% (range: 0.98%-8.89%) and 6.24% (range: 0.39%-19.04%) of the trials were rejected due to incorrect responses and artifacts respectively. Although ERP analyses were not the focus of the current study, they were performed for completion and references. The methods and detailed results of ERP analyses were reported in the Appendix.

### Multivariate Pattern Analysis (MVPA)

MVPA was implemented in MATLAB with custom-made codes and LIBSVM (Chang & Lin, 2011) using a linear support vector machine (c = 1). <u>The analysis code is publicly available on Figshare</u>. Classification was conducted in each participant separately with a sliding time window approach, using a window size of 50ms and a step size of 10ms. A subset of 64 posterior channels (Figure 3) was selected for the MVPA analysis based on the results of a pilot study, which showed that the general decoding accuracy was better with these channels than with all channels, regardless of conditions. We employed a ten-fold cross validation. The trials for classification were randomly assigned to ten bins of roughly equal number of trials and equal proportion of the two classes. Nine bins were pooled to train the classifier and the remaining one bin was used to test its classification accuracy. This step was repeated ten times so that every bin would serve as the test bin once. Principle Component Analysis (PCA) was applied to the 50 (time points) X 64 (channels) features in each time window in the training trials to reduce dimensionality. Principle components (PCs) explaining >99% variance were kept and used as input to train the classifier. The same PCs were applied to the testing trials when testing the classifiers’ performance. The purpose of conducting PCA on training trials only, instead of performing it on all trials combined, is to avoid spurious generalizability of training results to the testing data as a result of ’double-dipping’ in the feature selection procedure (Kriegeskorte et al., 2009; Mwangi et al., 2014). This cross validation procedure was repeated five times. The resulting accuracy of a classifier in a time window was the average accuracy of these five iterations of ten-fold cross validation.

**Figure 3.**
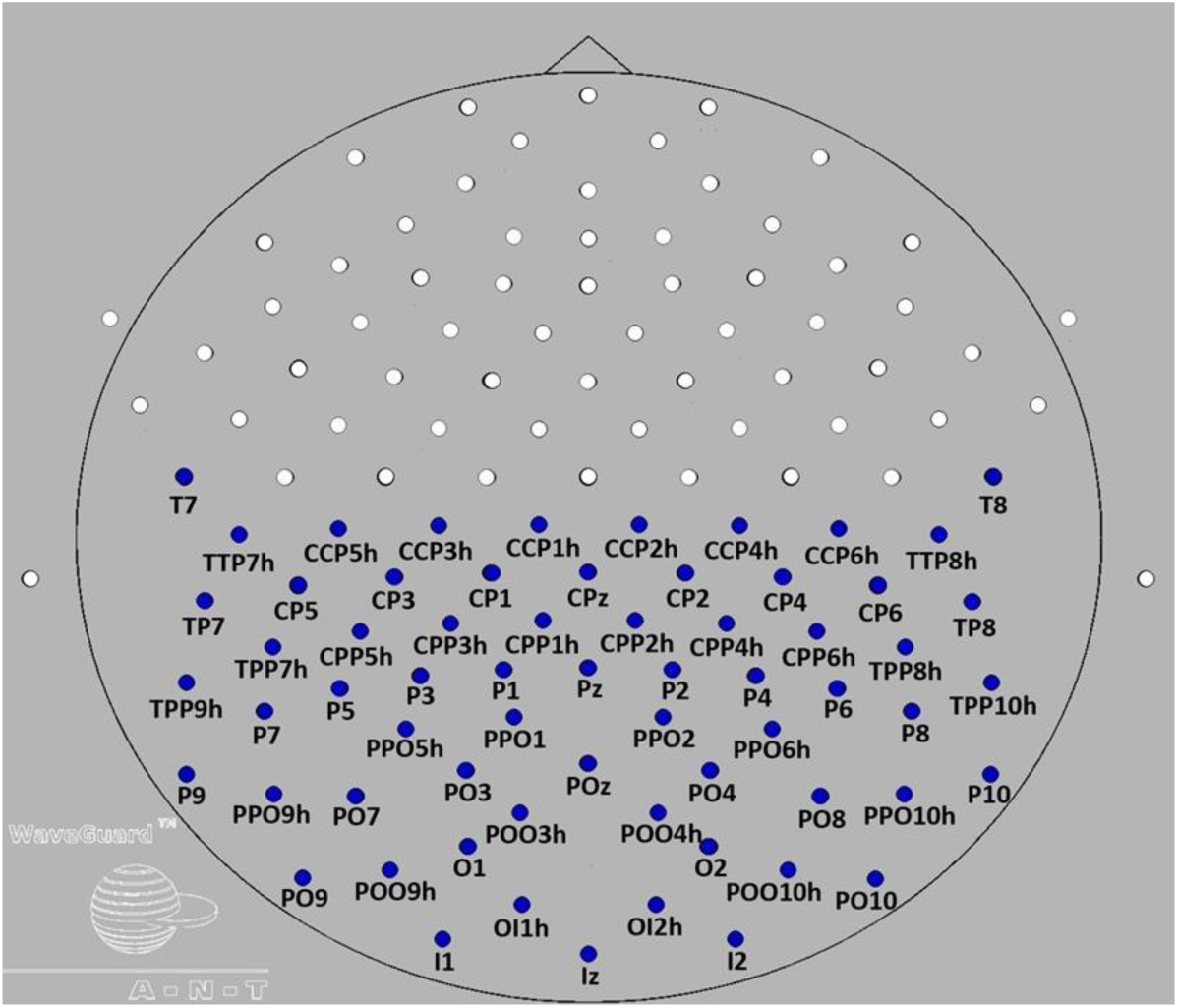
Channels used in MVPA were highlighted in blue.

Two major analyses were conducted to examine the effect of task on object processing. The first analysis involved comparison between the performance of task-relevant and task-irrelevant classifiers (Table 1). Task-relevant classifiers refer to the classifiers trained to discriminate an object dimension (e.g., texture) congruent with the participants’ task demand (e.g., as in a texture task trial). In contrast, task-irrelevant classifiers refer to the classifiers trained to discriminate an object dimension (e.g., texture) incongruent with the participants’ task demand (e.g., as in a shape task trial). If the task demand could render the object representations more separable along the dimension required by the task (e.g., a larger difference between the representations of Large-Round and Small-Round objects when participants were performing a texture vs. shape task), then the task-relevant classifiers would demonstrate higher average decoding accuracy than the task-irrelevant classifiers.

Another major analysis concerned whether the classifiers trained on one task can be generalized to another task. Performance of task-relevant and across-task classifiers was compared. Both types of classifiers were trained in the object dimension congruent with that of the participants’ task demand, however they were tested on data in different tasks. Task-relevant classifiers were tested on data in the same task with which the classifiers were trained, while across-task classifiers were tested on data in a different task from which the classifiers were trained, e.g., the Large-Round vs. Small-Round classifiers were trained on the texture task trials but tested on the shape task trials. If the features used for distinguishing the same objects differ between tasks, then what the classifiers learnt in one task would not be fully generalized to another task. This would result in a lower average decoding accuracy in the across-task classifiers than the task-relevant classifiers.

To perform significance testing on decoding accuracies, we used a non-parametric sign permutation test with a maximum cluster size method to correct for multiple comparisons (Nichols & Holmes, 2002; Hebart et al. 2018). First, we calculated a difference score at each time window for each participant. To test if the decoding accuracies of a classifier were above chance, we subtracted 0.5 from the accuracies. To test if the decoding accuracies of two classifiers were different, we subtracted the accuracies of one from another. Two-tailed tests with participants as the random factor were conducted to test whether the difference scores were significant. A permutation procedure was used in which each participant was randomly assigned a value of 1 or -1 and the difference scores at each time window was multiplied by this value. For each time window, a t-value was computed with the mean and SD of this score across participants. Repeating the procedure for 10,000 times, a distribution of 10,000 t-values was obtained for every time window. The primary threshold was set at the 2.5th and 97.5th percentile of this distribution. To obtain the secondary cluster-size threshold, we searched for clusters of significant time windows. Neighboring time windows would form a cluster only if all their t-values exceeded the primary threshold. The maximum cluster size of each permutation set was recorded, and it formed a distribution of maximum cluster sizes. The 97.5th percentile of this distribution served as a secondary cluster-size threshold. Finally, both thresholds were applied to the difference score of each time window. The difference score in a time window would be regarded as statistically significant if the t-value of the empirical data was higher than that of the primary threshold and if the size of the cluster encompassing this time window was larger than the secondary cluster-size threshold.

To estimate the effect size of differences in decoding, we computed a non-parametric estimator of common-language effect size measure *A* (Delaney & Vargha, 2002; Ruscio, 2008; Li, 2016) for the average accuracy across time within the significant clusters. It measures the probability of a randomly selected score from one condition being higher than a randomly selected score from another condition. We chose this measure because it requires no assumption on the underlying distribution as other measures like Cohen’s d (Cohen, 1988) do. The criteria of small, medium, and large effect size for common language measure are 0.56, 0.64, 0.71 respectively (Ruscio, 2008; Li, 2016). We reviewed our non-parametric measures with the same criteria. Apart from the effect size index, we also reported the percentage of individuals showing the same direction of effect as the group-averaged results to draw a more comprehensive picture.

## Results

### Behavioral Results

Overall responses were accurate (>93%) and fast (mean =570ms for correct trials). Figure 4 and 5 show the difference of accuracy and response time between task for each object group. We employed logit mixed model for the analysis of accuracy. We included task, object group and the interaction of task and object group as the fixed factors, and subject as a random intercept. Wald chi-square test of the model coefficients showed that main effect of task was not significant (χ^2^ = 1.82, df = 1, p = .178), while the main effect of object group (χ^2^ = 21.33, df = 3, p <.001) and interaction (χ^2^ = 11.28, df = 3, p =.01) were significant. Post hoc analysis of pairwise contrasts (Tukey-adjusted) showed that task difference was only evident in small-round objects (log odds ratio = -0.4502, z ratio = -4.435. p<.001).

**Figure 4.**
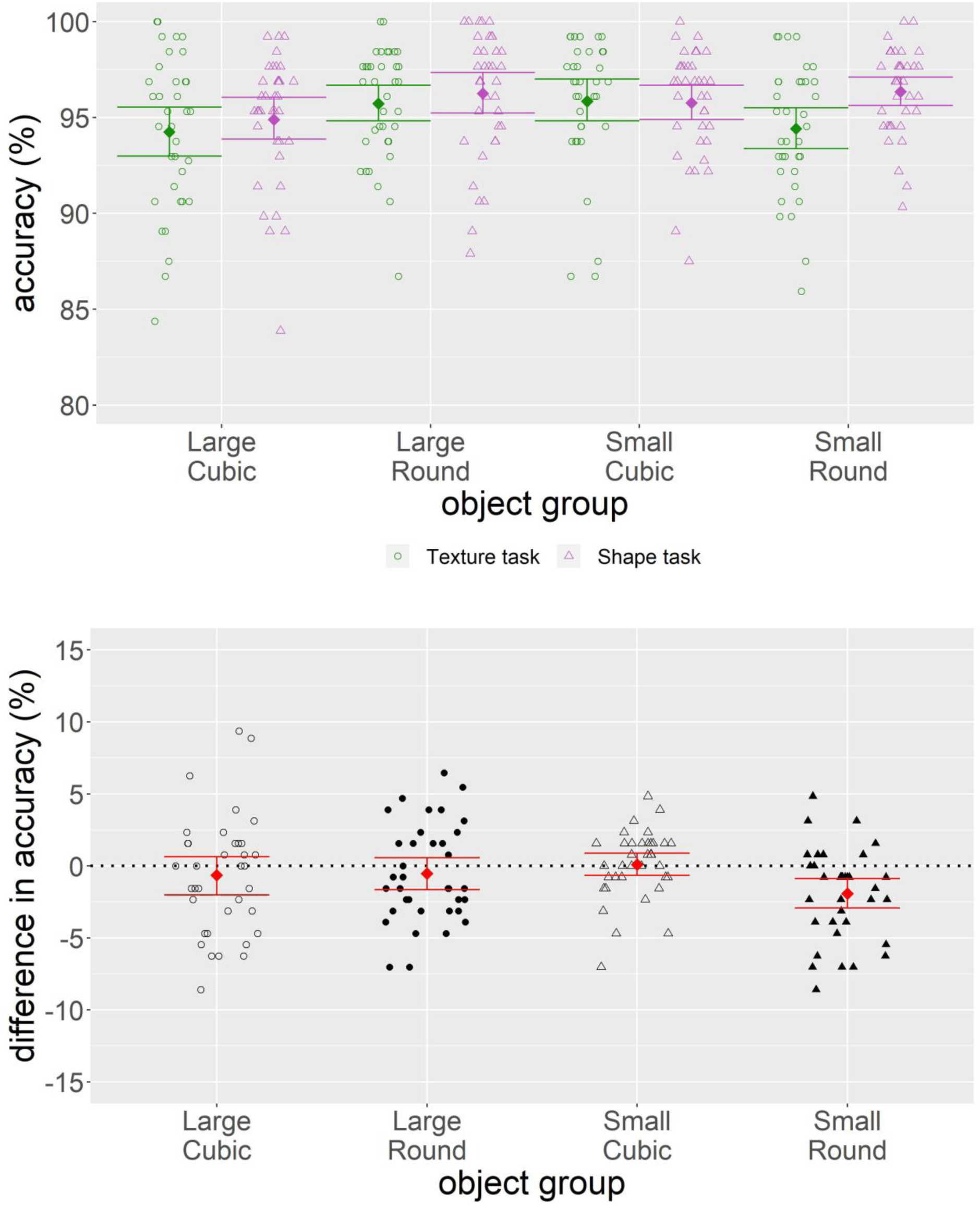
Behavioral accuracy of each object group in each task (top panel) and the difference between tasks (bottom panel). Each point represents one participant. Colored squares denote the mean across participants. Error bars represent 95% <u>percentile</u> bootstrap confidence interval (10000 samples) of the mean estimate.

**Figure 5.**
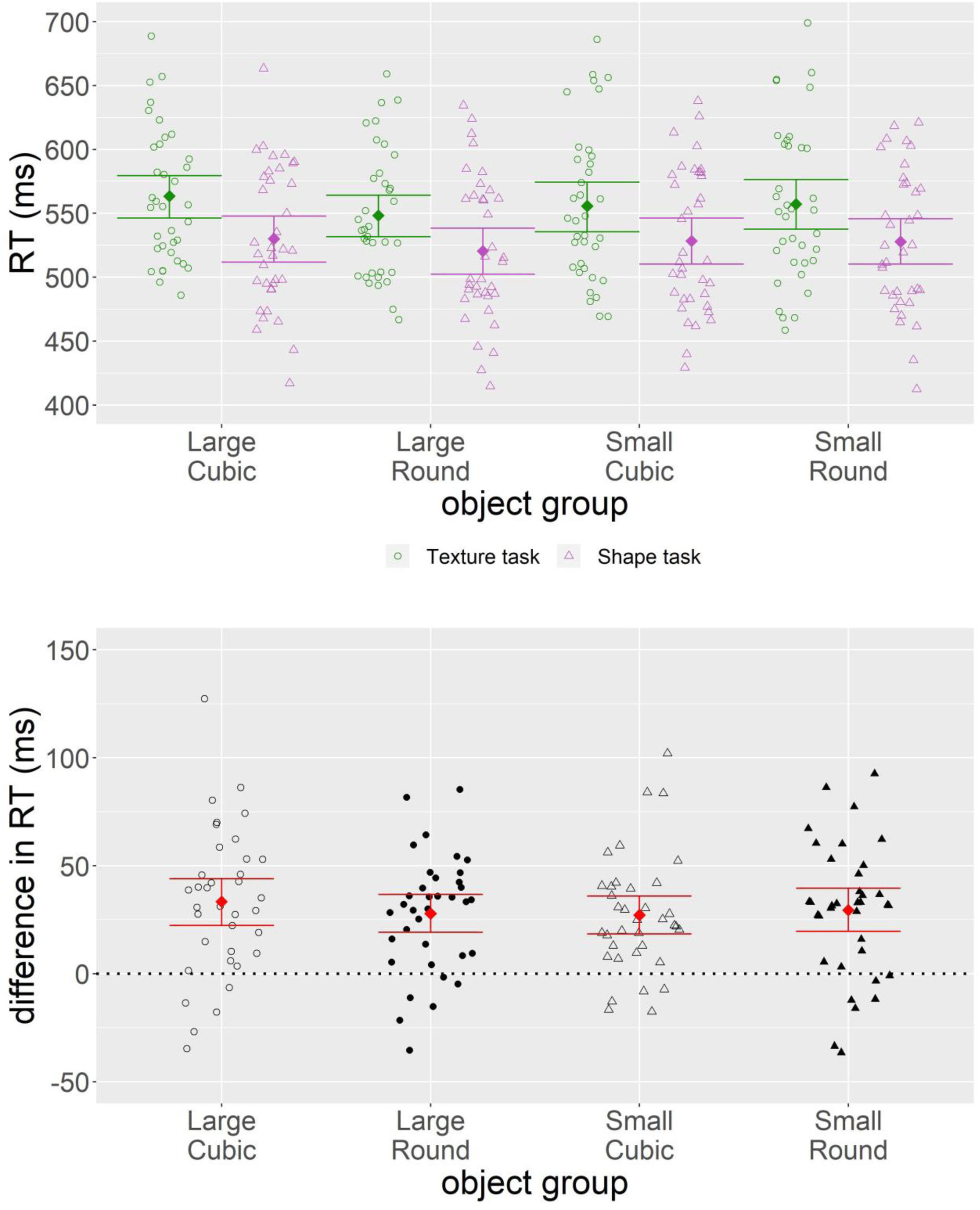
Mean response time across trials of each object group in each task (top panel) and the difference between tasks (bottom panel). Each point represents one participant. Colored squares denote the mean across participants. Error bars represent 95% <u>percentile</u> bootstrap confidence interval (10000 samples) of the mean estimate.

For response time data, we fit it with a linear mixed model, using task, object group and the interaction of task and object group as the fixed factors, and subject as a random intercept. Wald chi-square test of the model coefficients showed that all effects were significant, including task (χ^2^ = 103.02, df = 1, p = <.001), object group (χ^2^ = 49.28, df = 3, p <.001) and interaction (χ^2^ = 8.07, df = 3, p =.045). Post hoc analysis of pairwise contrasts (Tukey-adjusted) showed significant task difference in all object groups (estimates = 25.29 to 35.09, z ratio = 9.28 to 12.80, all p < .001).

### ERP Results

We first examined the difference between two tasks with all object groups collapsed. Figure 6 and 7 show the ERP waveforms and difference waveforms (texture task – shape task) in eight channels respectively. Figure 8 shows the scalp map of EEG signals in the two tasks across time. The selected channels showed clear deflections corresponding to the P1, the N170 and the N250 components. Repeated measure ANOVA analysis comparing the amplitudes of the N170 and the N250 in the two tasks showed that there were no significant difference in the N170 amplitudes (Figure 9), but the N250 amplitudes in the texture task trials were larger than that of shape task trials in the right hemisphere (Figure 10<u>). Details of the analysis and results were included in the Appendix.</u>

**Figure 6.**
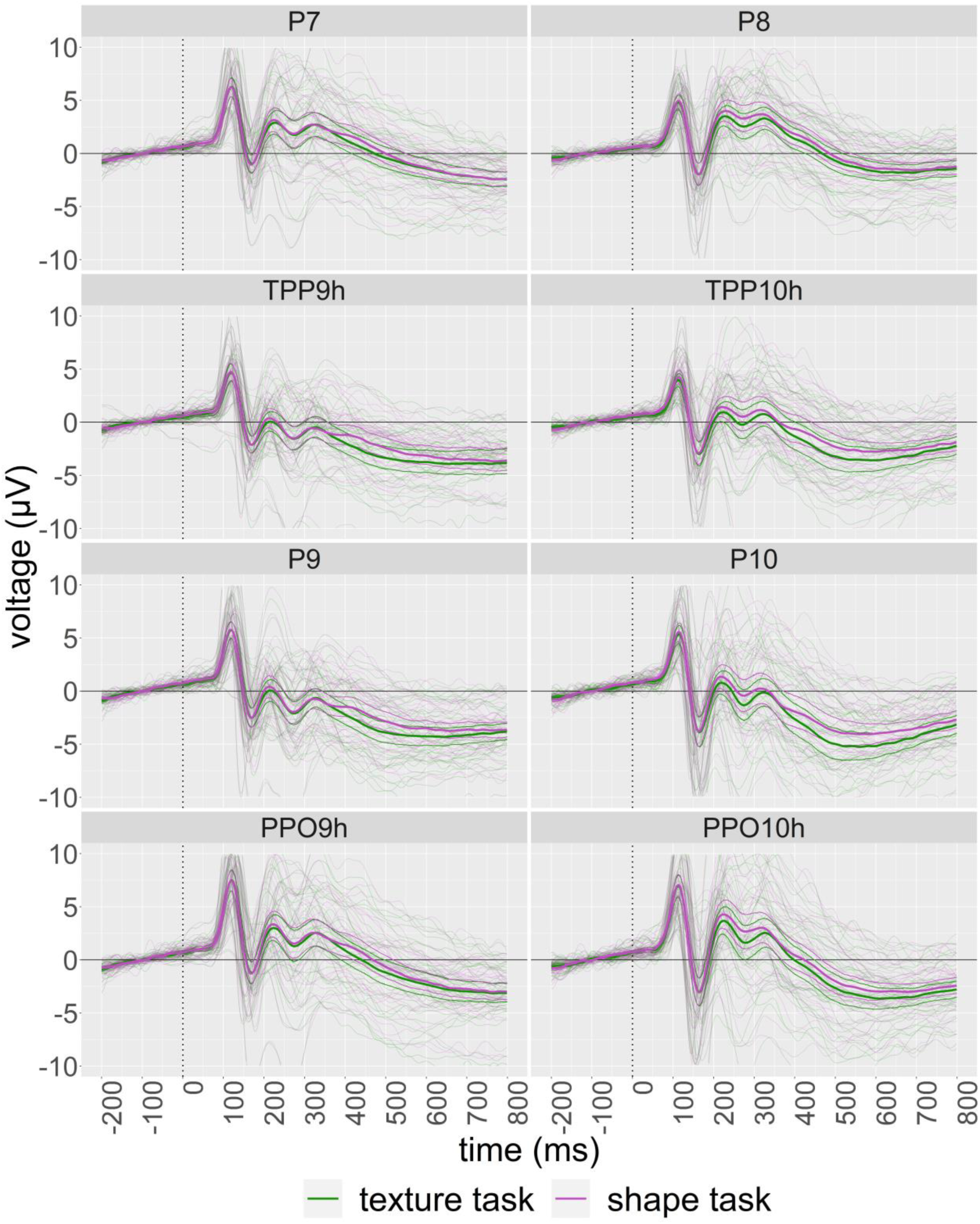
Grand average ERP waveform with individual plots superimposed in selected channels. Thinner curves around the grand average plot shows 95% <u>percentile</u> bootstrap confidence intervals (10000 samples) of the mean.

**Figure 7.**
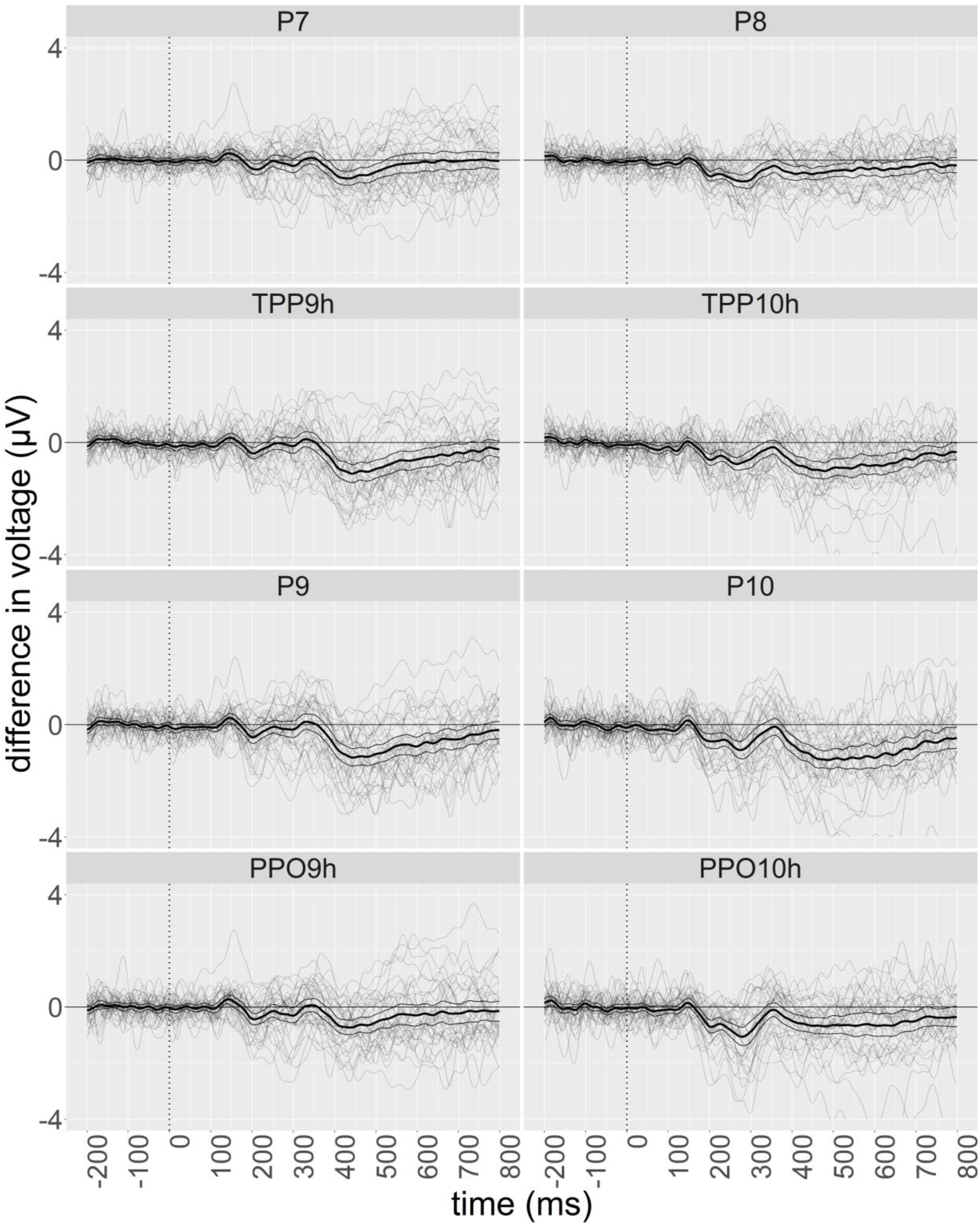
Grand average ERP difference waveform between task (texture minus shape) with individual plots superimposed in selected channels. Thinner curves around the grand average plot shows 95% <u>percentile</u> bootstrap confidence intervals (10000 samples) of the mean.

**Figure 8.**
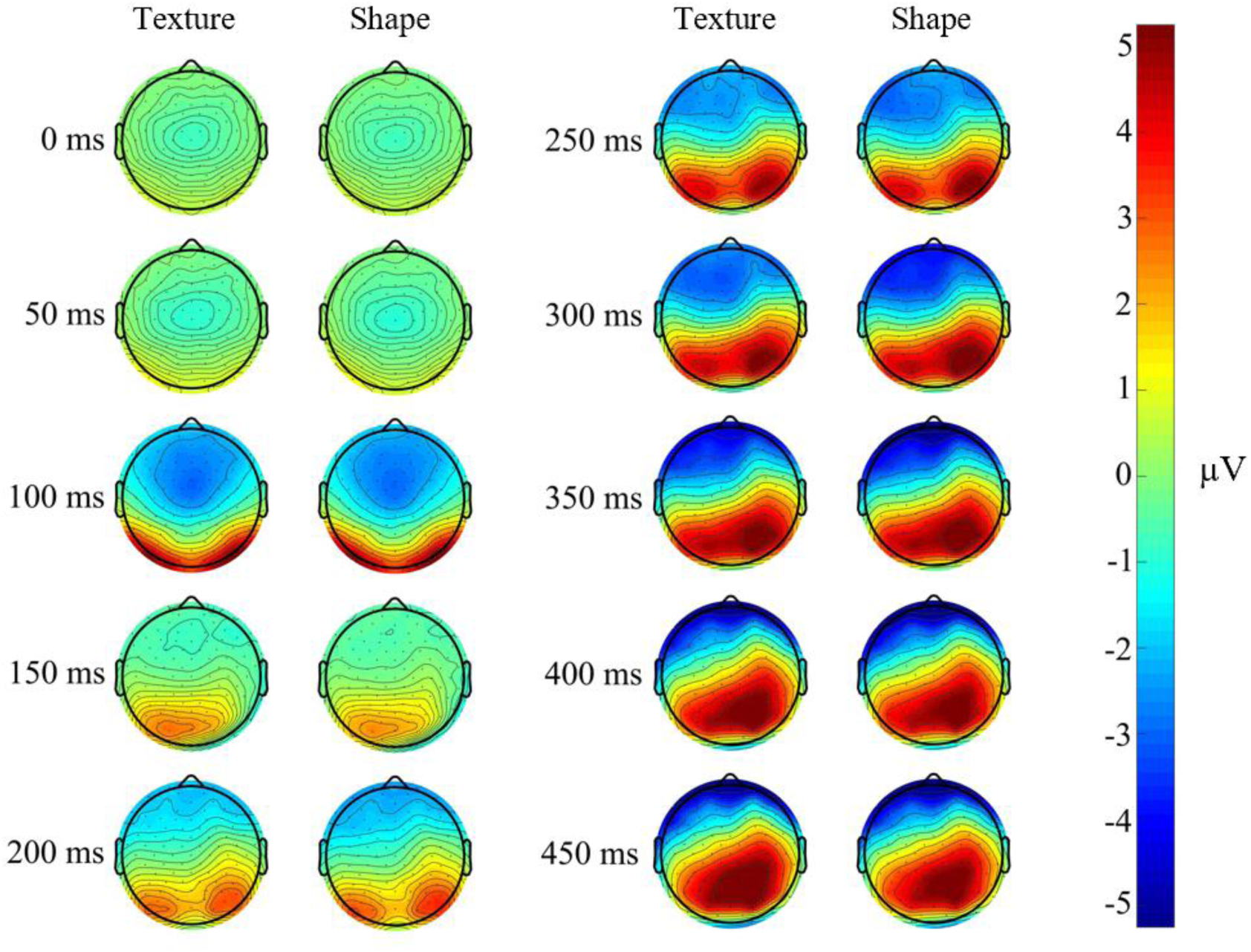
Scalp map of grand average ERP

**Figure 9.**
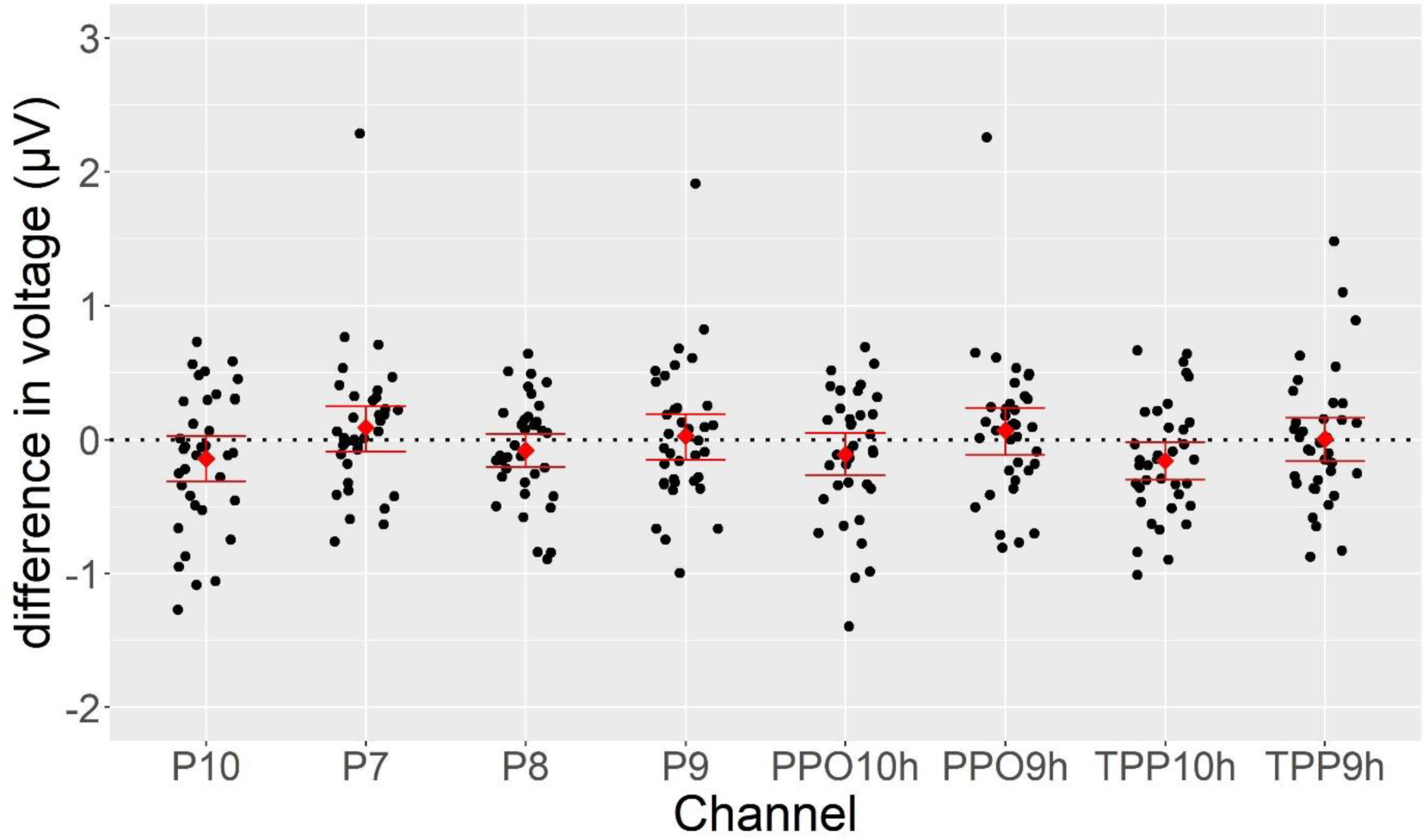
The N170 amplitude difference (texture - shape) in selected channels. Red squares represent mean estimate across participants. Error bars represent 95% percentile bootstrap confidence intervals (10000 samples) of the mean estimate.

**Figure 10.**
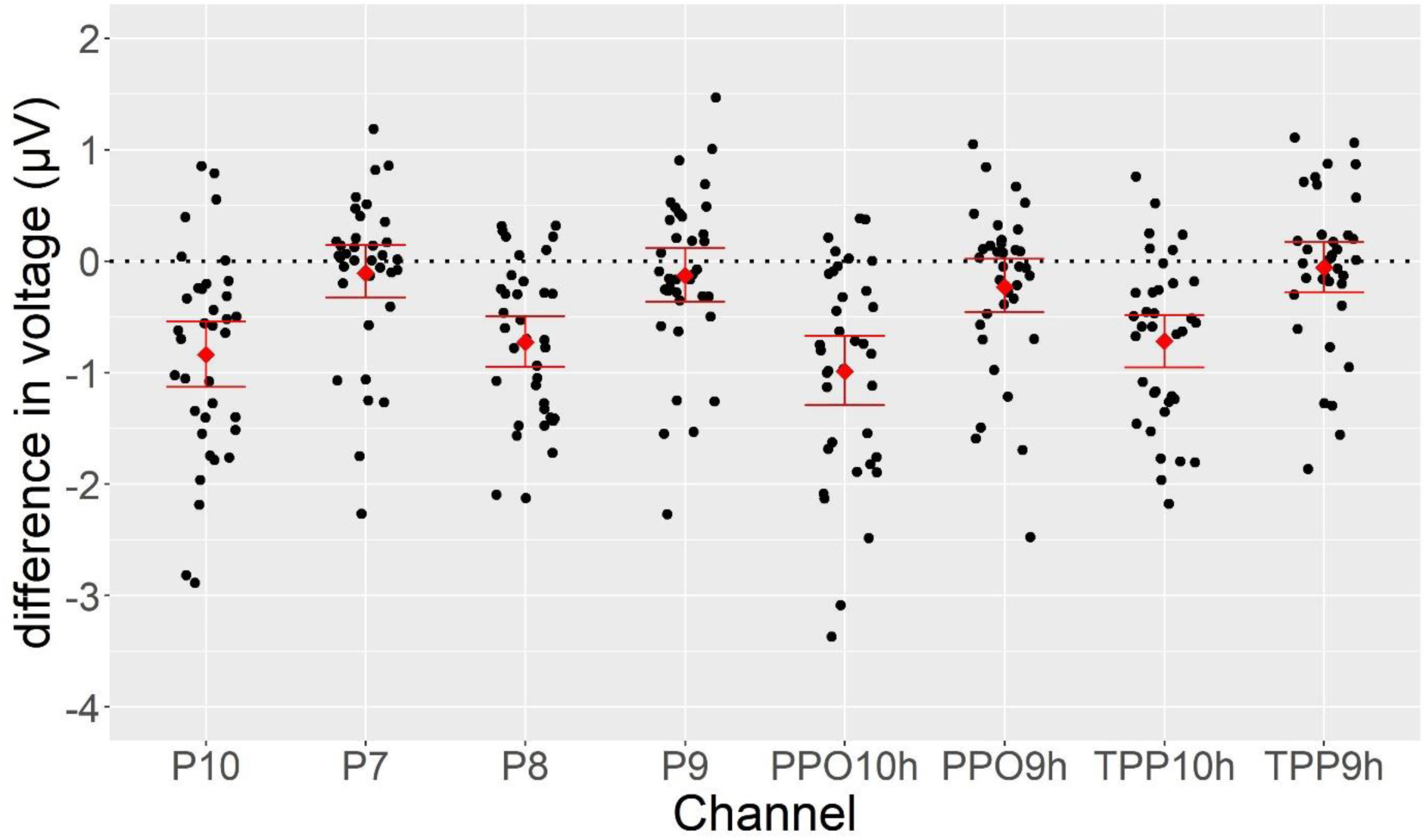
The N250 amplitude difference (texture - shape) in selected channels. Red squares represent mean estimate across participants. Error bars represent 95% percentile bootstrap confidence intervals (10000 samples) of the mean estimate.

We also computed the difference of ERP between large/small and round/cubic in the two tasks respectively (figures and details were included in the Appendix). For the large-small difference, we subtracted the average of ERP response of Small-Round and Small-Cubic from that of Large-Round and Large-Cubic. This difference was task-relevant in Texture task but task-irrelevant in Shape task. The round-cubic difference was computed in a similar way, and it is task-relevant in Shape task but task-irrelevant in Texture task. Repeated Measures ANOVA on the N170 and the N250 for both differences revealed no significant effect of task or interaction of task and channel.

### MVPA Results

Decoding accuracies of task-relevant, task-irrelevant and across-task classifiers (Table 1) across time were illustrated in Figure 11 (Shape classification) and Figure 12 (Texture classification). Overall, performance of both classifiers rose significantly above chance very early in time (70-130 in texture classification and 50-100ms in shape classification) and sustained significance until the end of epoch. For texture classification, peak latency ranged from 120-170ms to 140-190ms, and peak amplitudes were 60.21% (task-relevant), 59.22% (task-irrelevant) and 59.90% (across-task). For shape classification, peaks latency ranged from 130-180ms to 140-190ms, and peak amplitudes were 69.08% (task-relevant), 69.63% (task-irrelevant) and 68.74% (across-task).

**Figure 11.**
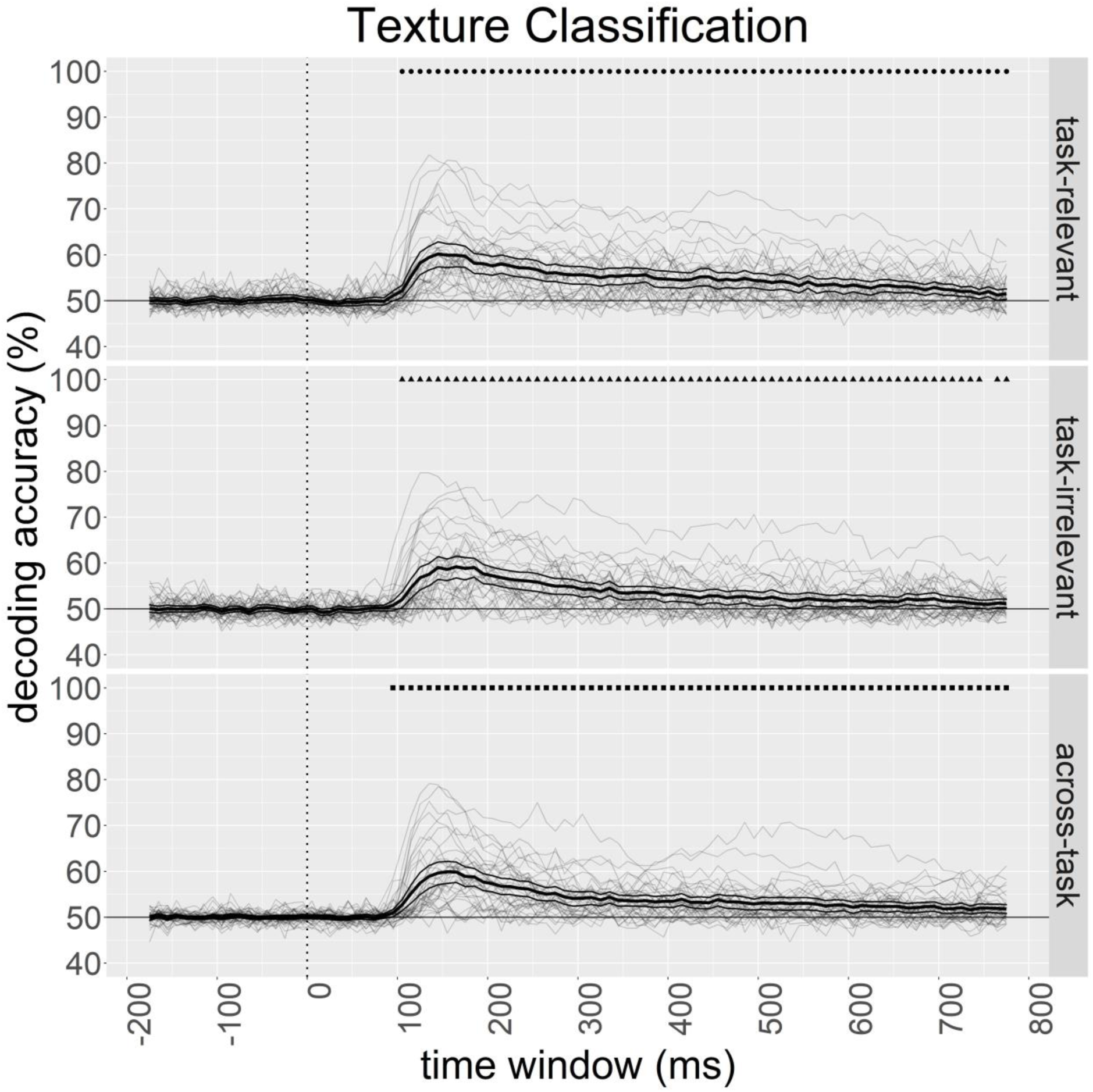
Mean decoding accuracies of task-relevant (top panel), task-irrelevant (middle panel) and across-task classifiers (bottom panel) for texture classification with individual plots superimposed. Thinner curves around the grand average plot shows 95% <u>percentile</u> bootstrap confidence intervals (10000 samples) of the mean. Dots at the top of the plots show time windows that the mean decoding accuracy was significantly different from chance (i.e. 50%).

**Figure 12.**
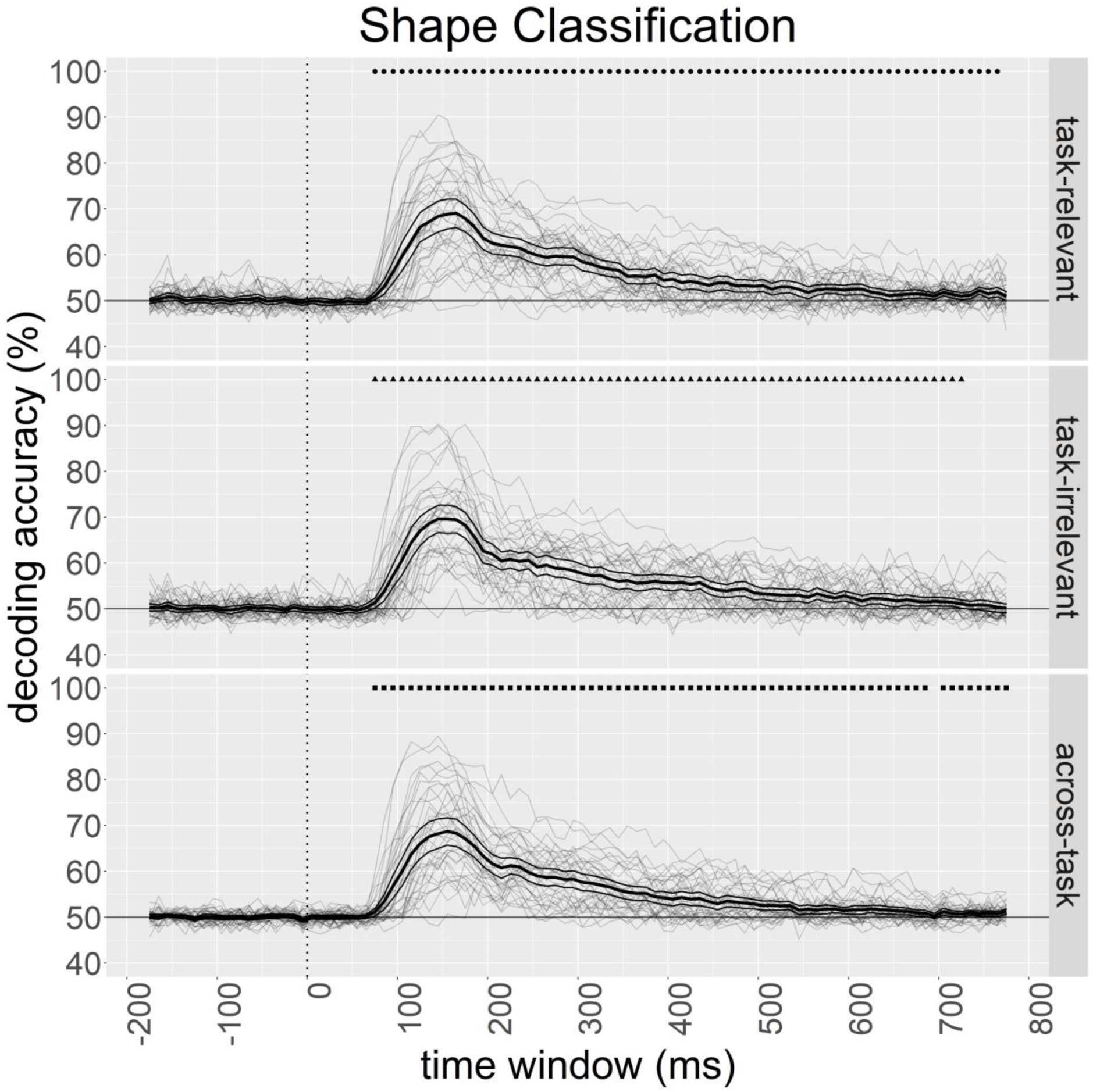
Mean decoding accuracies of task-relevant (top panel), task-irrelevant (middle panel) and across-task classifiers (bottom panel) for shape classification with individual plots superimposed. Thinner curves around the grand average plot shows 95% <u>percentile</u> bootstrap confidence intervals (10000 samples) of the mean. Dots at the top of the plots show time windows that the mean decoding accuracy was significantly different from chance (i.e. 50%).

In the first decoding analysis, performance of task-relevant classifiers was compared with that of task-irrelevant classifiers to test whether task relevance affected the amount of information useful for object differentiation. There were no significant differences between the decoding accuracy of task-relevant classifiers and that of task-irrelevant classifiers at all time windows in shape classification (top panel, Figure 14). In contrast, the comparison was significant in texture classification (top panel, Figure 13) at a relatively late time of 400-530ms (mean = 2.09%), suggesting higher amount of information was available for decoding in the task-relevant than task-irrelevant contexts. The effect size was medium-to-large, *A* = 0.672.

**Figure 13.**
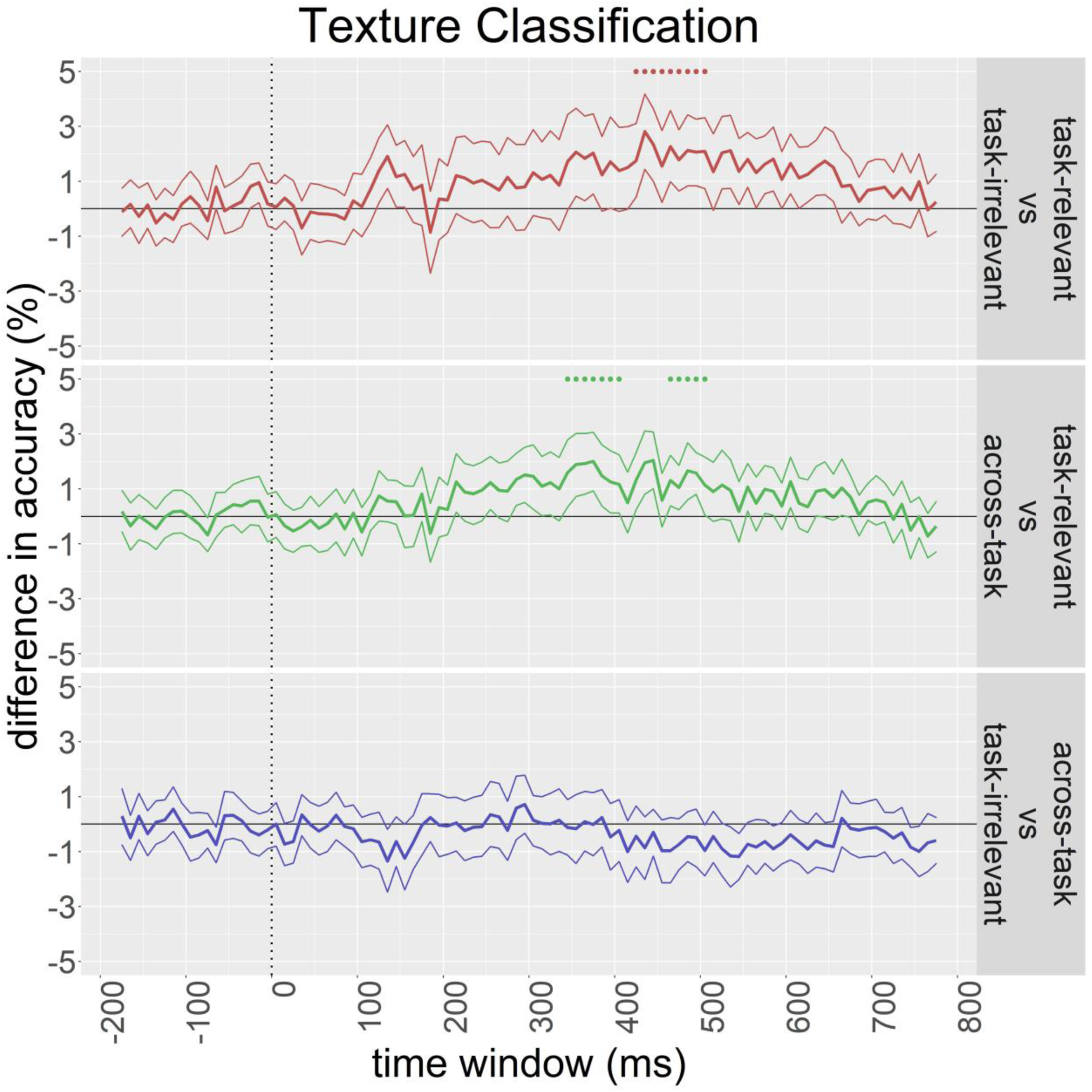
Mean difference of decoding accuracies in the comparison of task-relevant vs task-irrelevant classifiers (top panel), task-relevant vs across-task classifiers (middle panel), and task-irrelevant vs across-task classifiers (bottom panel) for texture classification. Thinner curves around the grand average plot shows 95% <u>percentile</u> bootstrap confidence intervals (10000 samples) of the mean. Dots at the top of the plots show time windows that the mean decoding accuracies of the two classifiers were significantly different. A similar plot with individual plots superimposed is included in the Appendix.

**Figure 14.**
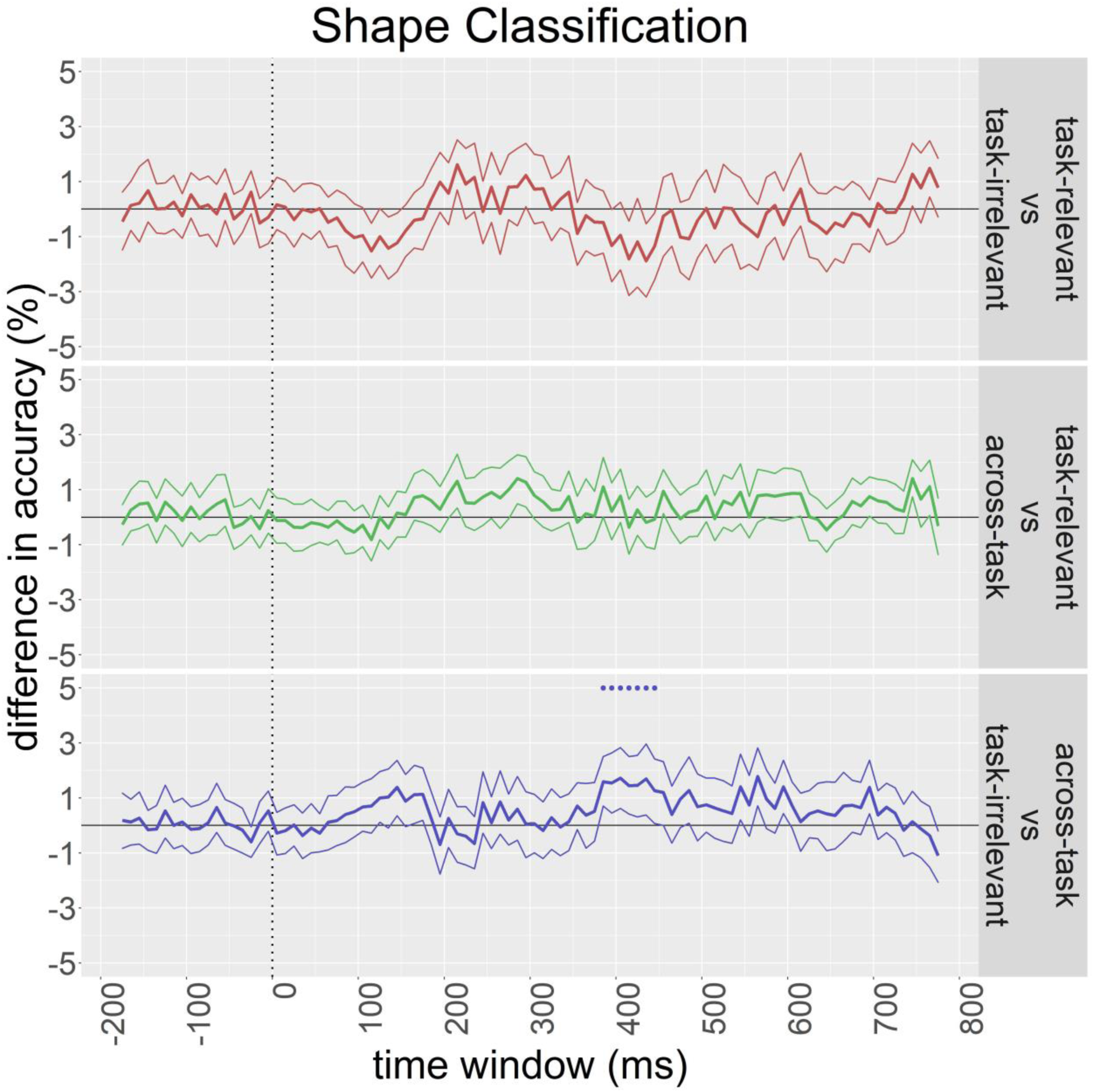
Mean difference of decoding accuracies in the comparison of task-relevant vs task-irrelevant classifiers (top panel), task-relevant vs across-task classifiers (middle panel), and task-irrelevant vs across-task classifiers (bottom panel) for shape classificationThinner curves around the grand average plot shows 95% <u>percentile</u> bootstrap confidence intervals (10000 samples) of the mean. Dots at the top of the plots show time windows that the mean decoding accuracies of the two classifiers were significantly different. A similar plot with individual plots superimposed is included in the Appendix.

In the second analysis, performance of task-relevant classifiers was compared with that of across-task classifiers to test whether task affected the source of information useful for object differentiation, or in other words, how well the features extracted by trained classifiers in one task generalized to a different task. There were no significant differences in the decoding accuracies of task-relevant classifiers and that of across-task classifiers across time in shape classification (middle panel, Figure 14. The same contrast, however, reached significance in texture decoding at 320-430ms (mean = 1.62%) and 440-530ms (mean = 1.35%; middle panel, Figure 13). It indicated that there was a lack of full generalization of what the classifiers learnt. The effect sizes of both clusters were small-to-medium, A = 0.588 and 0.593 respectively.

The third analysis compared task-irrelevant classifiers with across-task classifiers to test the unique information in across-task classifiers that was beyond task-irrelevant conditions. While both were tested in task-irrelevant conditions, across-task classifiers were trained in task-relevant conditions, whereas task-irrelevant classifiers were trained with task-irrelevant trials. Higher decoding accuracies in across-task classifiers would indicate extra information in task-relevant conditions that could help decoding in the task-irrelevant trials. The decoding accuracies of across-task classifiers were significantly higher at 360-470ms (mean = 1.53%) in shape classification (bottom panel, Figure 14) but remained non-significant for all time windows in texture classification (bottom panel, Figure 13). The effect size was small-to-medium, *A* = 0.610. It implied that the classifiers picked up some information in the task-relevant conditions that could better decode the task-irrelevant trials.

Figure 15 showed the averaged difference scores across time for each significant cluster. The percentage of participants showing the same direction as the group-averaged results ranged from 69.4% to 81%, suggesting that the majority of the participants showed the effects in a consistent direction.

**Figure 15.**
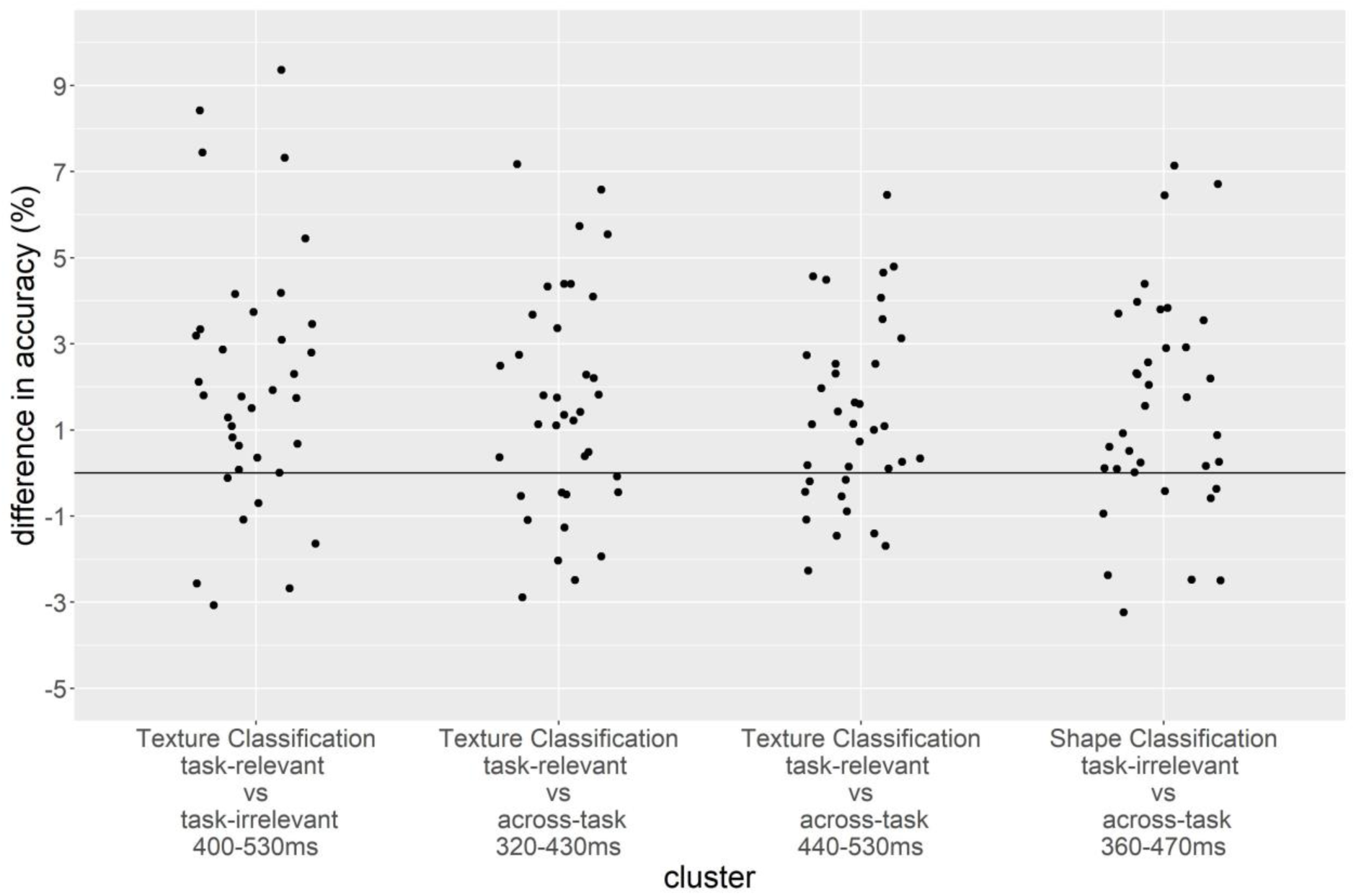
Average differences in decoding accuracy across time for each significant cluster. Each data point represents an individual participant. The percentages of participants that showed the same direction as the group-averaged results (i.e., difference > 0), from left to right, were 81%, 69.4%, 72.2% and 77.8%.

Given that our tasks required participants to attend to different feature dimensions, the frontal regions might potentially contribute to the differences in object processing between tasks. We thus performed decoding with frontal electrodes that were excluded in the above analyses. Overall, the decoding accuracies hovered above chance, with little significant difference between different classifiers. The data therefore did not contain information about the role of the frontal lobe in the task effect on object processing. Results were reported in Appendix.

## Discussion

The current study examined whether, when and how task affects object processing with decoding of EEG signals. We presented artificial stimuli that could be categorized along two dimensions (texture and shape) and asked participants to perform object judgment based on one dimension during each task.

### Task Effect on Object Representations

Overall, we found the source and amount of information for classification differed between tasks in texture classification. Specifically, we found a medium-to-large effect in the difference of decoding accuracies between task-relevant and task-irrelevant classifiers in texture classification at 400-530ms, suggesting that the amount of information in the brain activation pattern useful for object distinction along the texture dimension was higher in task-relevant than irrelevant contexts. Small-to-medium effects were found in the differences between task-relevant and across-task classifiers in texture classification at 320-430ms, and at 440-530ms. This suggests that task affected the features (channels, time points) that constitute to the information useful for object distinction along the texture dimension. In other words, there existed some qualitative differences in the object representations in the two tasks during this period, and the diagnostic features for object differentiation differed across task contexts.

Several implications arise from the finding of task effect on both the source and the amount of information in object representations. First, the findings may help resolve the inconsistent findings of the effect of task in previous studies. Studies that analyzed only the amount of information in brain activations and found no task effect might have overlooked the possibility that task may change the pattern of activations in certain situations. Here we showed that when the task demand changed, the channels and the time points that carry useful information did change accordingly, suggesting a recruitment of different sets of neural substrates for different task demands. By correlating representational dissimilarity matrix (RDM) of ERPs and S1 layer in HMAX model, Robinson et al. (2017) showed that the distinct patterns of activation across EEG channels in response to different spatial frequency represented the differences in neural activities in V1. Similarly, the lack of full generalization in the across-task decoding suggested that the pattern of activation, and therefore the underlying neural substrates, were distinguishable under different task demands. Second, the studies of object processing would likely benefit from the use of multiple types of tasks. One should beware that the object representations examined in a certain paradigm could change qualitatively in a different paradigm. Similarities and differences found between two object categories may therefore not hold when the task is changed. Finally, it is encouraging to see that EEG, with a lower spatial resolution than techniques like fMRI and MEG, is also capable of demonstrating different neural activation patterns across conditions. At least in the current context, multivariate analyses revealed dependence of object representations on task relevance that was not shown in univariate ERP analyses.

Across all participants task effects were evident in texture classification but not in shape classification. Given that participants were less accurate and had longer RT in the texture task, our results could be confounded with task difficulty: the larger task effect in the texture task might be solely due to participants paying more attention, effort etc., to this more difficult task. On the other hand, the visual processing required in the two tasks might be different. The shape task required more coarse processing while the texture task required more fine-level processing. Task difficulty and nature could therefore explain the finding of task effect in the texture classification but not shape classification. Future studies that matched task difficulty and precision levels of the required visual processing of the two tasks will help clarify to what degree these factors moderate the task effect on visual object processing.

Similar to the current study, Hebart et al. (2018) conducted analyses concerning both the amount of information contained in the activation patterns and the exact activation patterns across task contexts. However, they only found differences in the amount of information but not the qualitative difference of activation patterns between tasks. This discrepancy might be driven by the differences in the stimuli, tasks and analyses in the two studies. First, in our study, the objects used were novel, while Hebart et al. (2018) used a set of real-world objects, and due to our rich experience interacting with them for different purposes at the same time, it may have been difficult for observers to follow the instruction and focus only on dimensions relevant to the task demand. Second, in our study, both tasks involved were perceptual tasks, so they likely engaged partially different sets of neural substrates for perceptual analyses. In Hebart et al. (2018), the perceptual and conceptual tasks differed in depth of processing and could thus involve similar perceptual analyses and different subsequent analyses of conceptual information. Third, our study specifically examined the separability of object representations along different dimensions consistent and inconsistent with the task context, while in Hebart et al. (2018) classification was conducted for all objects in general without comparison between object classifications more consistent with the perceptual vs. conceptual task. Together this may have led to a task effect manifested in terms of the qualitative differences in the object representations in our study than in Hebart et al. (2018).

Effect size was not explicitly reported in Hebart et al. (2018), but we used the source data provided to calculate a measure of *A*. We averaged the accuracies within the significant cluster (542-833ms) of difference between perceptual and conceptual tasks and found that *A* = 0.803. It was larger than the effect we found in the comparison of task-relevant and task-irrelevant conditions in texture classification (*A* = 0.672). The distinction between the perceptual and conceptual tasks was the depth of processing, while that in the texture and shape tasks was the dimension of object categorization. A larger effect size in Hebart et al. (2018) could therefore be interpreted as a larger influence of the depth of processing by tasks than the dimension of categorization on object representation.

### Time-course of Task Effect

Task effect was found in several time windows starting from 320ms in texture classification, which lags behind the duration of the N170 and the N250. Consistent with the decoding results, our ERP difference (large-small and round-cubic) analysis showed no difference between tasks in the amplitudes of the N170 and the N250. EEG source localization studies have identified the fusiform gyrus as a potentially major source of the N170 (Deffke et al., 2007; Herrmann et al., 2005) and the N250 (Schweinberger et al., 2002; Schweinberger et al., 2004; Scott et al., 2006). Moreover, Cichy & Pantazis (2017) used fMRI-MEG/EEG fusion technique to study the representational structure of objects in time and space, and found correspondence between representations revealed by EEG decoding at different time points and representations revealed by fMRI at different areas along the ventral visual pathway. Specifically, representations at the primary visual cortex (V1) corresponded the best with MEG/EEG representations at about 100ms, while representations at the inferotemporal cortex (IT) corresponded the best with MEG/EEG representations at close to 300ms. Taken together, our results implied that task affects the late but not early stages of object processing.

Comparing to previous EEG/MEG decoding studies, Hebart et al. (2018) found differences in decoding accuracies of object categories in conceptual tasks and perceptual tasks starting between 542ms and 833ms, which was also relatively late. However, Bocincova & Johnson (2019) found that classifier performance in the decoding of a part’s orientation was higher when it was task-relevant than irrelevant during the 100-600ms time window. Therefore, the latency we found was somewhat in-between the two. One potential reason for the difference is that a blocked design was used in Bocincova & Johnson (2019) and participants were motivated to remember only the task-relevant feature in each block. In Hebart et al. (2018) and the current study, however, the task cue varied trial-by-trial. The frequent switch of task, and therefore the variable relevant feature required, made it hard for participants to focus on only particular features as in the blocked design. It may explain a later onset of task effect in Hebart et al. (2018) and our study. The block design may also explain the larger effect size observed in Bocincova & Johnson (2019) (η_p_^2^ = .397-.480) than that of the current study.

In conclusion, we found that object representations involved quantitatively and qualitatively different information across different task contexts. Task effects occurred after the time range of the N170 and the N250, which corresponds to later phases of object processing.

## Acknowledgements

We thank Prof. Maurer and Prof. Penney for their constructive comments. We also thank Kelvin Lui, Fan Pu and Daniel Yip for their assistance on the project.

## Abbreviations

EEG: Electroencephalogram
ERP: Event-Related Potential
IT: inferior temporal cortex
MEG: magnetoencephalography
MVPA: Multivariate Pattern Analysis
RDM: Representational Dissimilarity Matrix

## Authors’ contributions

Conceptualization: Hoi Ming Ken Yip, Yetta Kwailing Wong, Alan C.-N. Wong

Data curation: Hoi Ming Ken Yip, Leo Y. T. Cheung

Formal Analysis: Hoi Ming Ken Yip, Yetta Kwailing Wong, Vince S. H. Ngan, Alan C.-N. Wong

Investigation: Hoi Ming Ken Yip, Leo Y. T. Cheung, Alan C.-N. Wong

Methodology: Hoi Ming Ken Yip, Leo Y. T. Cheung, Yetta Kwailing Wong, Vince S. H. Ngan, Alan C.-N. Wong

Project Administration: Hoi Ming Ken Yip, Leo Y. T. Cheung, Yetta Kwailing Wong, Alan C.-N. Wong

Resources: Leo Y. T. Cheung, Yetta Kwailing Wong, Alan C.-N. Wong

Software: Hoi Ming Ken Yip, Leo Y. T. Cheung, Yetta Kwailing Wong, Vince S. H. Ngan, Alan C.-N. Wong

Visualization: Hoi Ming Ken Yip, Yetta Kwailing Wong, Vince S. H. Ngan, Alan C.-N. Wong Supervision: Yetta Kwailing Wong, Alan C.-N. Wong

Writing original draft: Hoi Ming Ken Yip, Alan C.-N. Wong

Writing-review and editing: Hoi Ming Ken Yip, Leo Y. T. Cheung, Yetta Kwailing Wong, Vince S. H. Ngan, Alan C.-N. Wong

## References

Alilović, J., Timmermans, B., Reteig, L. C., Van Gaal, S., & Slagter, H. A. (2019). No evidence that predictions and attention modulate the first feedforward sweep of cortical information processing. Cerebral Cortex, 29(5), 2261–2278. https://doi.org/10.1093/cercor/bhz038

Bentin, S., Allison, T., Puce, A., Perez, E., & McCarthy, G. (1996). Electrophysiological studies of face perception in humans. Journal of cognitive neuroscience, 8(6), 551–565. https://doi.org/10.1162/jocn.1996.8.6.551

Bocincova, A., & Johnson, J. S. (2019). The time course of encoding and maintenance of task-relevant versus irrelevant object features in working memory. Cortex, 111, 196–209. https://doi.org/10.1016/j.cortex.2018.10.013

Bracci, S., Daniels, N., & Op de Beeck, H. (2017). Task context overrules object-and category-related representational content in the human parietal cortex. Cerebral Cortex, 27(1), 310–321. https://doi.org/10.1093/cercor/bhw419

Brainard, D. H. (1997). The psychophysics toolbox. Spatial vision, 10, 433–436. https://doi.org/10.1163/156856897x00357

Chang, C. C., & Lin, C. J. (2011). LIBSVM: A library for support vector machines. ACM transactions on intelligent systems and technology (TIST), 2(3), 27. https://doi.org/10.1145/1961189.1961199

Cichy, R. M., & Pantazis, D. (2017). Multivariate pattern analysis of MEG and EEG: A comparison of representational structure in time and space. NeuroImage, 158, 441–454. https://doi.org/10.1016/j.neuroimage.2017.07.023

Clark, A. (2013). Whatever next? Predictive brains, situated agents, and the future of cognitive science. Behavioral and brain sciences, 36(3), 181–204. https://doi.org/10.1017/s0140525x12000477

Cohen, J. (1988). Statistical power analysis for the behavioral sciences (2nd ed.). Hillsdale, NJ: Erlbaum.

Cohen Kadosh, K., Henson, R. N., Cohen Kadosh, R., Johnson, M. H., & Dick, F. (2010). Task-dependent activation of face-sensitive cortex: an fMRI adaptation study. Journal of Cognitive Neuroscience, 22(5), 903–917. https://doi.org/10.1162/jocn.2009.21224

Deffke, I., Sander, T., Heidenreich, J., Sommer, W., Curio, G., Trahms, L., & Lueschow, A. (2007). MEG/EEG sources of the 170-ms response to faces are co-localized in the fusiform gyrus. Neuroimage, 35(4), 1495–1501. https://doi.org/10.1016/j.neuroimage.2007.01.034

Delaney, H. D., & Vargha, A. (2002). Comparing several robust tests of stochastic equality with ordinally scaled variables and small to moderate sized samples. Psychological Methods, 7(4), 485. https://doi.org/10.1037/1082-989X.7.4.485

Delorme, A., & Makeig, S. (2004). EEGLAB: an open source toolbox for analysis of single-trial EEG dynamics including independent component analysis. Journal of neuroscience methods, 134(1), 9–21. https://doi.org/10.1016/j.jneumeth.2003.10.009

DiCarlo, J. J., Zoccolan, D., & Rust, N. C. (2012). How does the brain solve visual object recognition? Neuron, 73(3), 415–434. https://doi.org/10.1016/j.neuron.2012.01.010

Firestone, C., & Scholl, B. J. (2016). Cognition does not affect perception: Evaluating the evidence for “top-down” effects. Behavioral and brain sciences, 39. https://doi.org/10.1017/s0140525x15000965

Friston, K. (2010). The free-energy principle: a unified brain theory? Nature reviews neuroscience, 11(2), 127. https://doi.org/10.1038/nrn2787

Gauthier, I., & Tarr, M. J. (1997). Becoming a “Greeble” expert: Exploring mechanisms for face recognition. Vision research, 37(12), 1673–1682. https://doi.org/10.1016/s0042-6989(96)00286-6

Goffaux, V., Jemel, B., Jacques, C., Rossion, B., & Schyns, P. G. (2003). ERP evidence for task modulations on face perceptual processing at different spatial scales. Cognitive Science, 27(2), 313–325. https://doi.org/10.1207/s15516709cog2702_8

Grill-Spector, K., Kourtzi, Z., & Kanwisher, N. (2001). The lateral occipital complex and its role in object recognition. Vision research, 41(10-11), 1409–1422. https://doi.org/10.1016/s0042-6989(01)00073-6

Grootswagers, T., Wardle, S. G., & Carlson, T. A. (2017). Decoding dynamic brain patterns from evoked responses: A tutorial on multivariate pattern analysis applied to time series neuroimaging data. Journal of cognitive neuroscience, 29(4), 677–697. https://doi.org/10.1162/jocn_a_01068

Harel, A., Kravitz, D. J., & Baker, C. I. (2014). Task context impacts visual object processing differentially across the cortex. Proceedings of the National Academy of Sciences, 201312567. https://doi.org/10.1073/pnas.1312567111

Haynes, J. D. (2015). A primer on pattern-based approaches to fMRI: principles, pitfalls, and perspectives. Neuron, 87(2), 257–270. https://doi.org/10.1016/j.neuron.2015.05.025

Hebart, M. N., Bankson, B. B., Harel, A., Baker, C. I., & Cichy, R. M. (2018). The representational dynamics of task and object processing in humans. Elife, 7, e32816. https://doi.org/10.7554/elife.32816

Herrmann, M. J., Ehlis, A. C., Muehlberger, A., & Fallgatter, A. J. (2005). Source localization of early stages of face processing. Brain topography, 18(2), 77–85. https://doi.org/10.1007/s10548-005-0277-7

Hohwy, J. (2013). The predictive mind. Oxford University Press. https://doi.org/10.1093/acprof:oso/9780199682737.001.0001

Hohwy, J. (2017). Priors in perception: Top-down modulation, Bayesian perceptual learning rate, and prediction error minimization. Consciousness and Cognition, 47, 75–85. https://doi.org/10.1016/j.concog.2016.09.004 https://doi.org/10.1016/j.concog.2016.05.003

Itier, R. J., & Neath-Tavares, K. N. (2017). Effects of task demands on the early neural processing of fearful and happy facial expressions. Brain research, 1663, 38–50. https://doi.org/10.1016/j.brainres.2017.03.013

Jackson, J., Rich, A. N., Williams, M. A., & Woolgar, A. (2017). Feature-selective attention in frontoparietal cortex: multivoxel codes adjust to prioritize task-relevant information. Journal of cognitive neuroscience, 29(2), 310–321. https://doi.org/10.1162/jocn_a_01039

Jehee, J. F., Brady, D. K., & Tong, F. (2011). Attention improves encoding of task-relevant features in the human visual cortex. Journal of Neuroscience, 31(22), 8210–8219. https://doi.org/10.1523/jneurosci.6153-09.2011

Julian, J. B., Ryan, J., & Epstein, R. A. (2016). Coding of object size and object category in human visual cortex. Cerebral Cortex, 27(6), 3095–3109. https://doi.org/10.1093/cercor/bhw150

Kim, N. Y., & McCarthy, G. (2016). Task influences pattern discriminability for faces and bodies in ventral occipitotemporal cortex. Social neuroscience, 11(6), 627–636. https://doi.org/10.1080/17470919.2015.1131194

Kok, P., Jehee, J. F., & De Lange, F. P. (2012). Less is more: expectation sharpens representations in the primary visual cortex. Neuron, 75(2), 265–270. https://doi.org/10.1016/j.neuron.2012.04.034

Kriegeskorte, N., Mur, M., Ruff, D. A., Kiani, R., Bodurka, J., Esteky, H., … & Bandettini, P. A. (2008). Matching categorical object representations in inferior temporal cortex of man and monkey. Neuron, 60(6), 1126–1141. https://doi.org/10.1016/j.neuron.2008.10.043

Kriegeskorte, N., Simmons, W. K., Bellgowan, P. S., & Baker, C. I. (2009). Circular analysis in systems neuroscience: the dangers of double dipping. Nature Neuroscience, 12(5), 535–540.https://doi.org/10.1038/nn.2303

Li, J. C. H. (2016). Effect size measures in a two-independent-samples case with nonnormal and nonhomogeneous data. Behavior research methods, 48(4), 1560–1574. https://doi.org/10.3758/s13428-015-0667-z

Long, N. M., & Kuhl, B. A. (2018). Bottom-up and top-down factors differentially influence stimulus representations across large-scale attentional networks. Journal of Neuroscience, 2724-17. https://doi.org/10.1523/jneurosci.2724-17.2018

Lopez-Calderon, J., & Luck, S. J. (2014). ERPLAB: an open-source toolbox for the analysis of event-related potentials. Frontiers in human neuroscience, 8, 213. https://doi.org/10.3389/fnhum.2014.00213

Lupyan, G. (2015). Cognitive penetrability of perception in the age of prediction: Predictive systems are penetrable systems. Review of philosophy and psychology, 6(4), 547–569. https://doi.org/10.1007/s13164-015-0253-4

Macpherson, F. (2017). The relationship between cognitive penetration and predictive coding. Consciousness and cognition, 47, 6–16. https://doi.org/10.1016/j.concog.2016.04.001

Mwangi, B., Tian, T. S., & Soares, J. C. (2014). A review of feature reduction techniques in neuroimaging. Neuroinformatics, 12(2), 229–244. https://doi.org/10.1007/s12021-013-9204-3

Newen, A., & Vetter, P. (2017). Why cognitive penetration of our perceptual experience is still the most plausible account. Consciousness and cognition, 47, 26–37. https://doi.org/10.1016/j.concog.2016.09.005

Nichols, T. E., & Holmes, A. P. (2002). Nonparametric permutation tests for functional neuroimaging: a primer with examples. Human brain mapping, 15(1), 1–25. https://doi.org/10.1002/hbm.1058

O’Callaghan, C., Kveraga, K., Shine, J. M., Adams Jr, R. B., & Bar, M. (2017). Predictions penetrate perception: Converging insights from brain, behaviour and disorder. Consciousness and cognition, 47, 63–74.

Pelli, D. G. (1997). The VideoToolbox software for visual psychophysics: Transforming numbers into movies. Spatial vision, 10, 437–442. https://doi.org/10.1163/156856897x00366

Pylyshyn, Z. (1999). Is vision continuous with cognition?: The case for cognitive impenetrability of visual perception. Behavioral and brain sciences, 22(3), 341–365. https://doi.org/10.1017/s0140525x99002022

Raftopoulos, A. (2014). The cognitive impenetrability of the content of early vision is a necessary and sufficient condition for purely nonconceptual content. Philosophical Psychology, 27(5), 601–620. https://doi.org/10.1080/09515089.2012.729486

Ransom, M., Fazelpour, S., & Mole, C. (2017). Attention in the predictive mind. Consciousness and cognition, 47, 99–112. https://doi.org/10.1016/j.concog.2016.06.011

Riesenhuber, M., & Poggio, T. (2002). Neural mechanisms of object recognition. Current opinion in neurobiology, 12(2), 162–168. https://doi.org/10.1016/s0959-4388(02)00304-5

Robinson, A. K., Venkatesh, P., Boring, M. J., Tarr, M. J., Grover, P., & Behrmann, M. (2017). Very high density EEG elucidates spatiotemporal aspects of early visual processing. Scientific reports, 7(1), 1–11. https://doi.org/10.1038/s41598-017-16377-3

Ruscio, J. (2008). A probability-based measure of effect size: robustness to base rates and other factors. Psychological methods, 13(1), 19. https://doi.org/10.1037/1082-989X.13.1.19

Schwarzkopf, D. S., & Rees, G. (2011). Pattern classification using functional magnetic resonance imaging. Wiley Interdisciplinary Reviews: Cognitive Science, 2(5), 568–579. https://doi.org/10.1002/wcs.141

Schweinberger, S. R., Huddy, V., & Burton, A. M. (2004). N250r: a face-selective brain response to stimulus repetitions. Neuroreport, 15(9), 1501–1505. https://doi.org/10.1097/01.wnr.0000131675.00319.42

Schweinberger, S. R., Pickering, E. C., Burton, A. M., & Kaufmann, J. M. (2002). Human brain potential correlates of repetition priming in face and name recognition. Neuropsychologia, 40(12), 2057–2073. https://doi.org/10.1016/s0028-3932(02)00050-7

Scott, L. S., Tanaka, J. W., Sheinberg, D. L., & Curran, T. (2006). A reevaluation of the electrophysiological correlates of expert object processing. Journal of cognitive neuroscience, 18(9), 1453–1465. https://doi.org/10.1162/jocn.2006.18.9.1453

Serre, T., Kouh, M., Cadieu, C., Knoblich, U., Kreiman, G., & Poggio, T. (2005). A theory of object recognition: computations and circuits in the feedforward path of the ventral stream in primate visual cortex. MASSACHUSETTS INST OF TECH CAMBRIDGE MA CENTER FOR BIOLOGICAL AND COMPUTATIONAL LEARNING.

Serre, T., Oliva, A., & Poggio, T. (2007). A feedforward architecture accounts for rapid categorization. Proceedings of the national academy of sciences, 104(15), 6424–6429. https://doi.org/10.1073/pnas.0700622104

Tanaka, J. W., & Curran, T. (2001). A neural basis for expert object recognition. Psychological science, 12(1), 43–47. https://doi.org/10.1111/1467-9280.00308

Teufel, C., & Nanay, B. (2017). How to (and how not to) think about top-down influences on visual perception. Consciousness and cognition, 47, 17–25. https://doi.org/10.1016/j.concog.2016.05.008

Valdés-Conroy, B., Aguado, L., Fernández-Cahill, M., Romero-Ferreiro, V., & Diéguez-Risco, T. (2014). Following the time course of face gender and expression processing: a task-dependent ERP study. International Journal of Psychophysiology, 92(2), 59–66. https://doi.org/10.1016/j.ijpsycho.2014.02.005

Vance, J., & Stokes, D. (2017). Noise, uncertainty, and interest: Predictive coding and cognitive penetration. Consciousness and cognition, 47, 86–98. https://doi.org/10.1016/j.concog.2016.06.007

Vaziri-Pashkam, M., & Xu, Y. (2017). Goal-directed visual processing differentially impacts human ventral and dorsal visual representations. Journal of Neuroscience, 37(36), 8767–8782. https://doi.org/10.1523/jneurosci.3392-16.2017

Wong, A. C. N., Palmeri, T. J., & Gauthier, I. (2009). Conditions for facelike expertise with objects: Becoming a Ziggerin expert—but which type? Psychological Science, 20(9), 1108–1117. https://doi.org/10.1111/j.1467-9280.2009.02430.x

Wong, A. C., Gauthier, I., Woroch, B., Debuse, C., & Curran, T. (2005). An early electrophysiological response associated with expertise in letter perception. Cognitive, Affective, & Behavioral Neuroscience, 5(3), 306–318. https://doi.org/10.3758/cabn.5.3.306

