## appendix for "The Effect of Task on Object Processing revealed by EEG decoding"

### ERP Analysis Method

We compared the amplitudes of two ERP components, the N170 and N250, between conditions. For the N170, the channels were selected by examining the amplitude of the negative peak between 140ms-200ms in the grand average ERP across all participants and with all conditions collapsed. The channels with the largest amplitudes in the left and right hemisphere, P9 and P10, and six adjacent channels (P7, P8, TPP9h, TPP10h, PPO9h, PPO10h) were selected. Furthermore, the latencies of the negative peaks between 140ms-200ms in P9 and P10 in the same grand average ERP were identified and averaged. The averaged latency was 167ms. We defined the N170 as the mean amplitude within a 40ms time window centering at this latency, which was 147ms-187ms. The same eight channels were selected for the analysis of the N250. The latencies of the negative peak between 230-330ms in P9 and P10 in the same grand average ERP were identified and averaged. The resulting latency was 274ms, which defined a 40ms time window of 254ms-294ms for calculating the mean amplitude. Both the amplitudes of the N170 and the N250 were calculated for the texture task trials and shape task trials separately in the eight channels in each participant.

### Comparing ERP in two task contexts

A 2 (task: texture and shape) X 8 (channel: P7, P8, P9, P10, TPP9h, TPP10h, PPO9h, PPO10h) repeated measure ANOVA was performed on the amplitudes of the N170 and the

N250 respectively. For the N170 (Figure 8), the main effect of channel was significant, $F(7,29) = 13.252$, $p < .001$, $\eta_p^2 = .762$. There was no significant main effect of task, $F(1,35) = .319$, $p = .576$, $\eta_p^2 = .009$, or interaction between task and channel, $F(7,29) = 1.584$, $p = .180$, $\eta_p^2 = .277$.

For the N250 (Figure 9), there was significant main effect of task, $F(1,35) = 21.987$, $p < .001$, $\eta_p^2 = .386$, significant main effect of channels, $F(7,29) = 26.078$, $p < .001$, $\eta_p^2 = .863$, and significant interaction of task and channels, $F(7,29) = 4.842$, $p = .001$, $\eta_p^2 = .539$. Subsequent paired t-tests performed for each channel separately showed significantly more negative N250 amplitudes in the texture task than the shape task for the channels in the right hemisphere including P8, $t(35) = -6.231$, $p < .001$; P10, $t(35) = -5.839$, $p < .001$; TPP10h, $t(35) = -5.513$, $p < .001$ and PPO10h, $t(35) = -6.232$, $p<.001$ but not for the channels in the left hemisphere, $p>.05$.

**Comparing ERP differences in two task contexts**

We computed two ERP difference waveforms that resemble task-relevant and task-irrelevant analyses. The first one was large-small difference (waveform in Figure A1, scalp map in Figure A2), in which we subtracted the average of ERP response to Small-Round and Small-Cubic from that of Large-Round and Large-Cubic. Another one was round-cubic difference, which is the difference between Large-Round/Small-Round and Large-Cubic/Small-Cubic. We conducted two repeated measure ANOVA with the same factors as above in the N170 and the N250. For the analysis of the N170 in large-small difference (Figure A3), the main effect of channel was significant, $F(7,29) = 4.269$, $p =.002$, $\eta_p^2 = .507$. There was no significant main effect of task, $F(1,35) = .311$, $p = .581$, $\eta_p^2 = .009$, or interaction between task and channel, $F(7,29) = .288$, $p = .953$, $\eta_p^2 = .065$. For the N250 (Figure A4), there were no significant main effect of channel, $F(7,29) = 2.149$, $p =.07$, $\eta_p^2 = .342$, main effect of task, $F(1,35) = 1.031$, $p = .317$, $\eta_p^2 = .029$, or interaction between task and channel, $F(7,29) = .745$, $p = .636$, $\eta_p^2 = .152$.

Result patterns were similar in round-cubic difference (waveform in Figure A5, scalp map in Figure A6). The analysis of the N170 (Figure A7) revealed significant effect of channel, $F(7,29) = 5.450$, $p <.001$, $\eta_p^2 = .568$, but not main effect of task, $F(1,35) = .056$, $p = .815$, $\eta_p^2 = .002$, or interaction between task and channel, $F(7,29) = .576$, $p = .769$, $\eta_p^2 = .122$. For the N250 (Figure A8), there were no significant main effect of channel, $F(7,29) = 1.204$, $p =.332$, $\eta_p^2 = .225$, main effect of task, $F(1,35) = .004$, $p = .949$, $\eta_p^2 = .000$, or interaction between task and channel, $F(7,29) = 1.364$, $p = .258$, $\eta_p^2 = .248$.

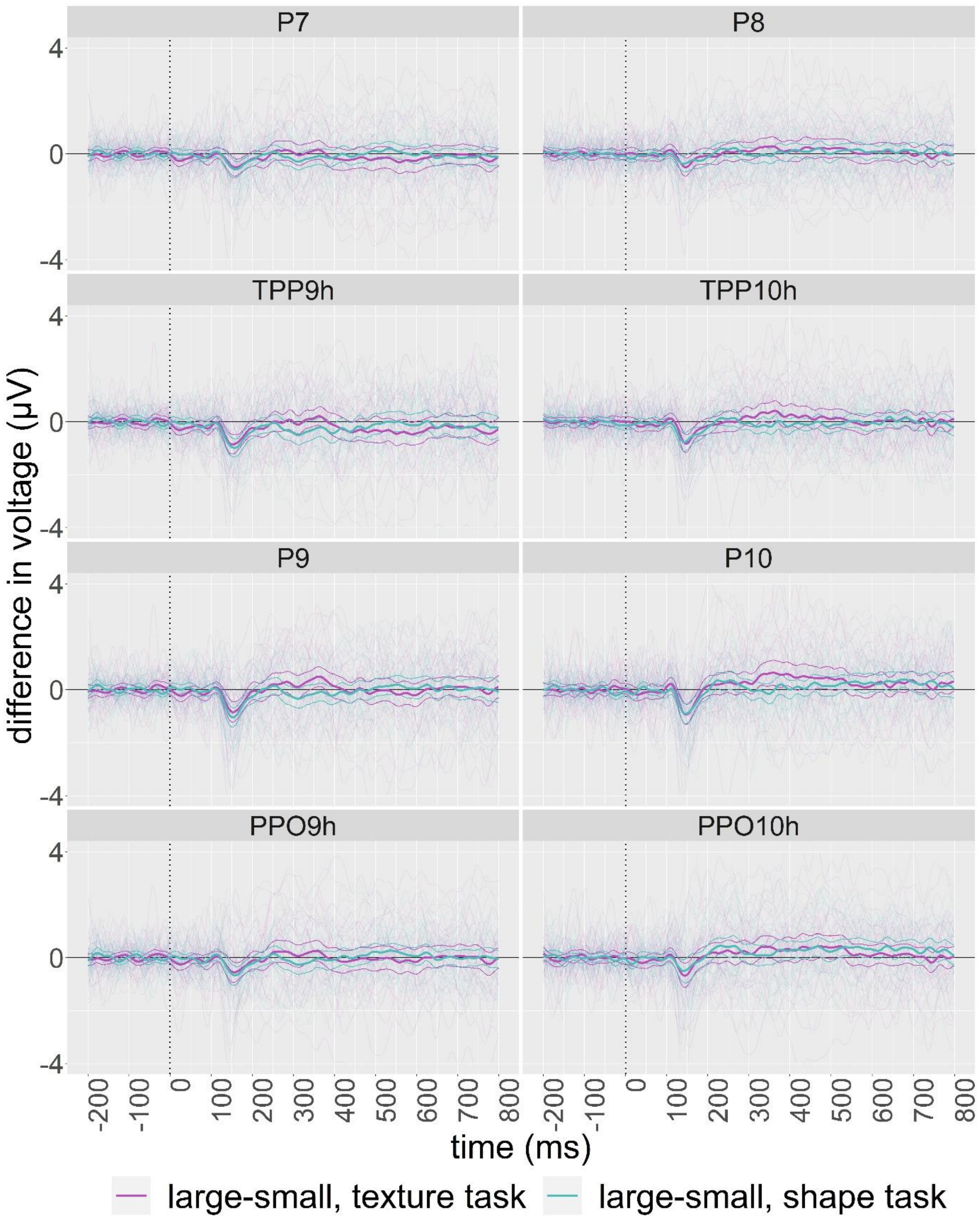


*Figure A1*. Grand average large-small difference ERP waveform with individual plots superimposed in selected channels. Thinner curves around the grand average plot shows 95% percentile bootstrap confidence intervals (10000 samples) of the mean.

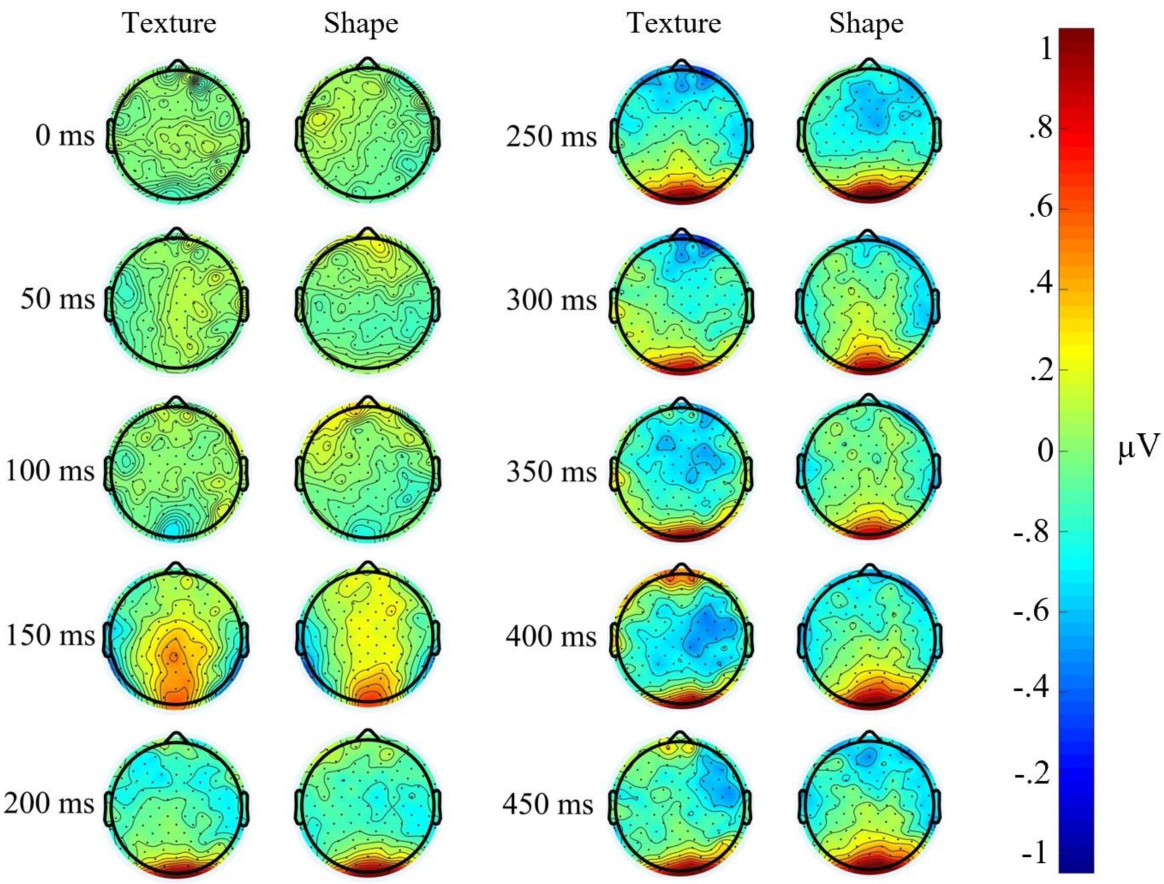


*Figure A2.* Scalp map of grand average large-small difference ERP

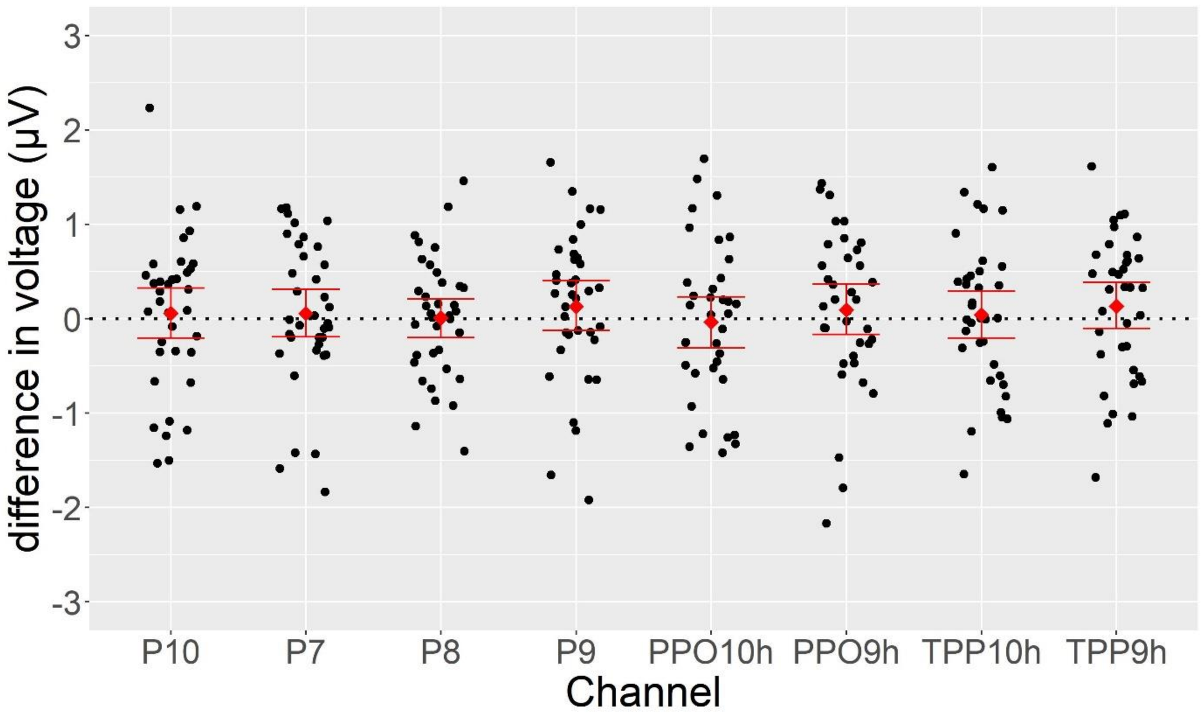


*Figure A3*. The N170 amplitude difference (texture - shape) in large-small difference in selected channels. Red squares represent mean estimate across participants. Error bars represent 95% percentile bootstrap confidence intervals (10000 samples) of the mean estimate.

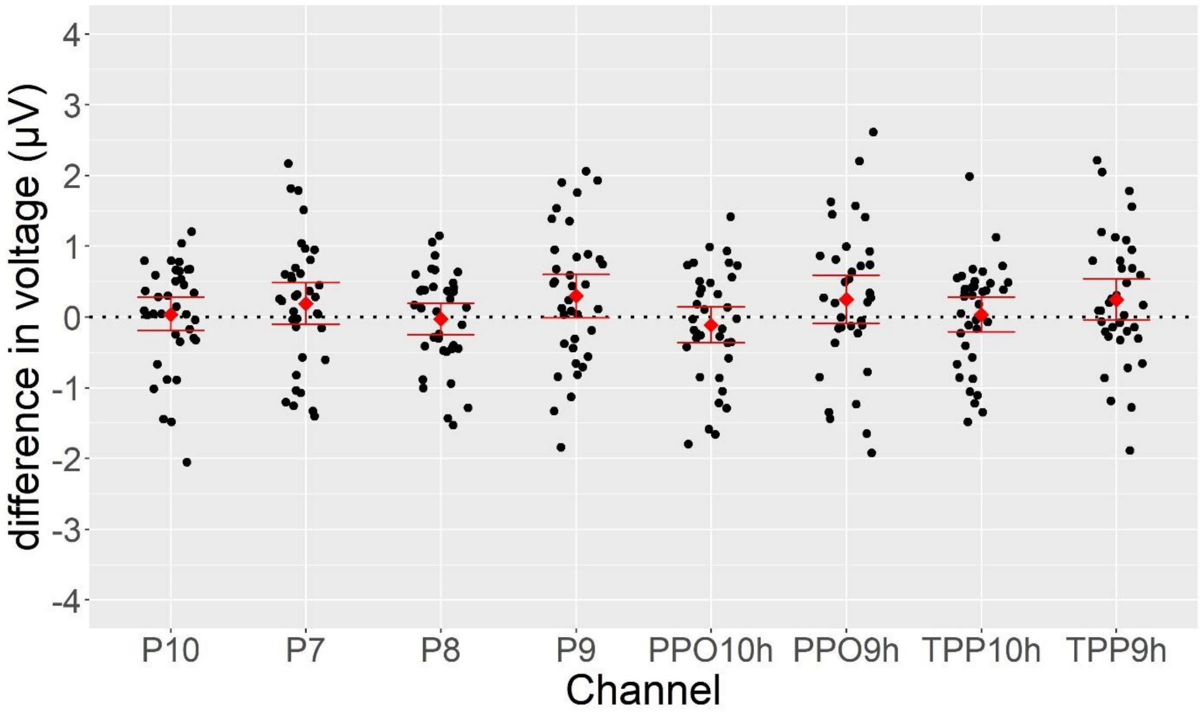


*Figure A4*. The N250 amplitude difference (texture - shape) in large-small difference in selected channels. Red squares represent mean estimate across participants. Error bars represent 95% percentile bootstrap confidence intervals (10000 samples) of the mean estimate.

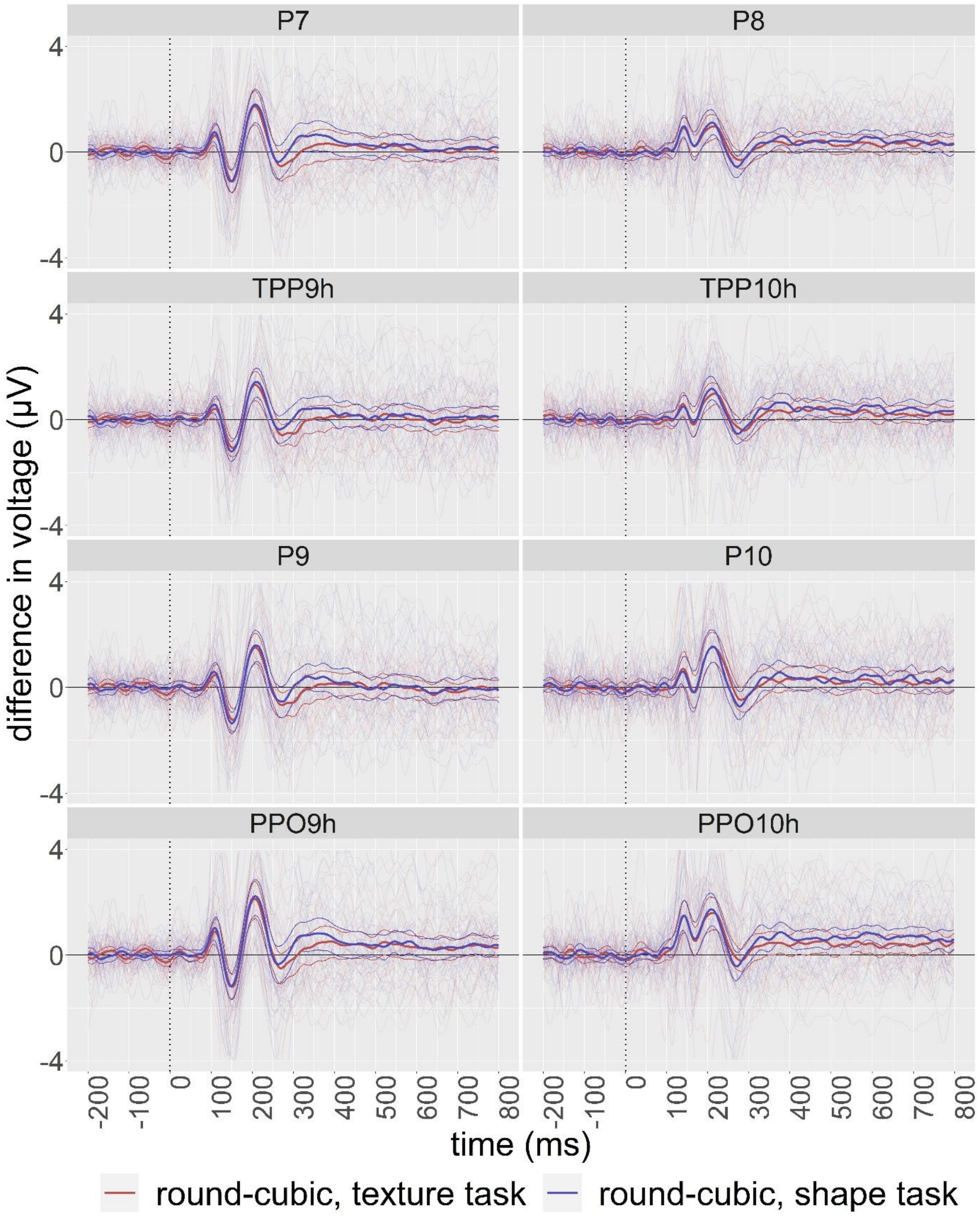

*Figure A5.* Grand average round-cubic difference ERP waveform with individual plots superimposed in selected channels. Thinner curves around the grand average plot shows 95% percentile bootstrap confidence intervals (10000 samples) of the mean.

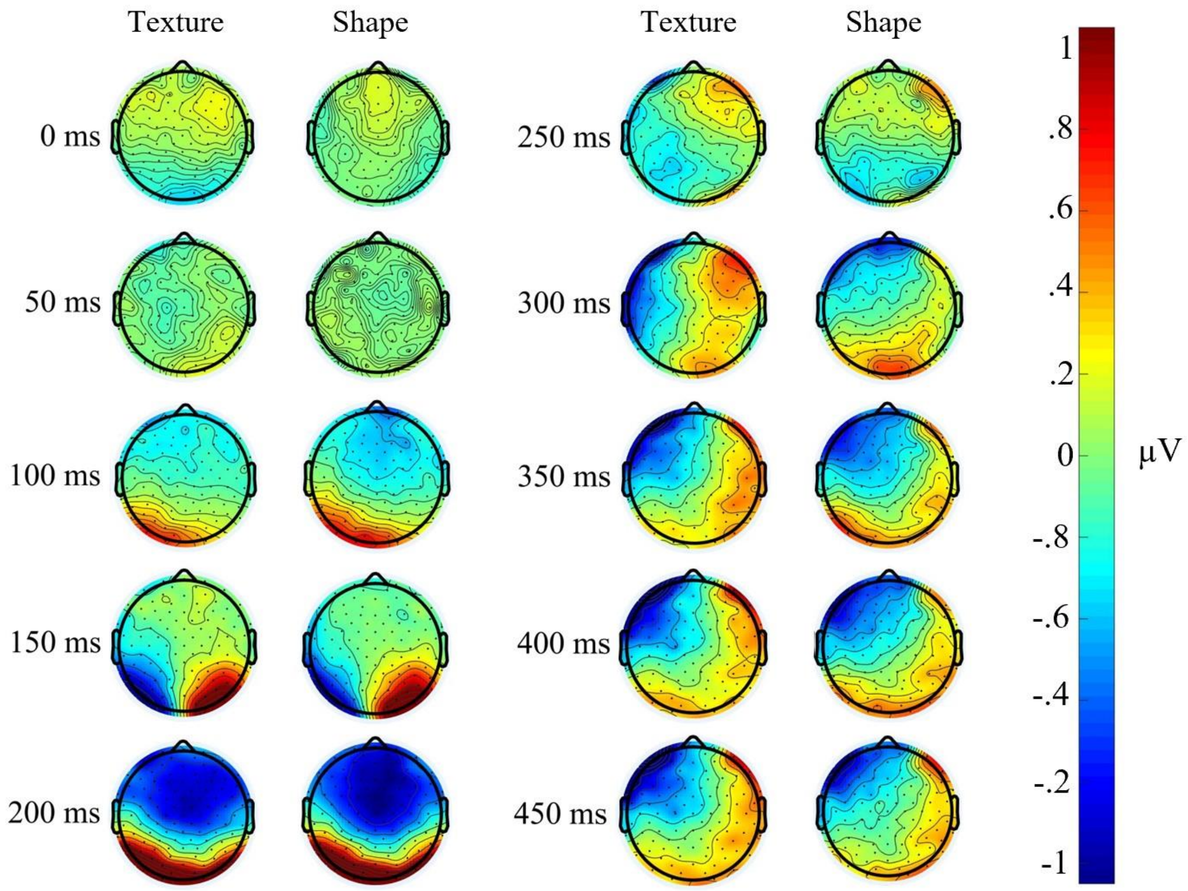


*Figure A6.* Scalp map of grand average round-cubic difference ERP

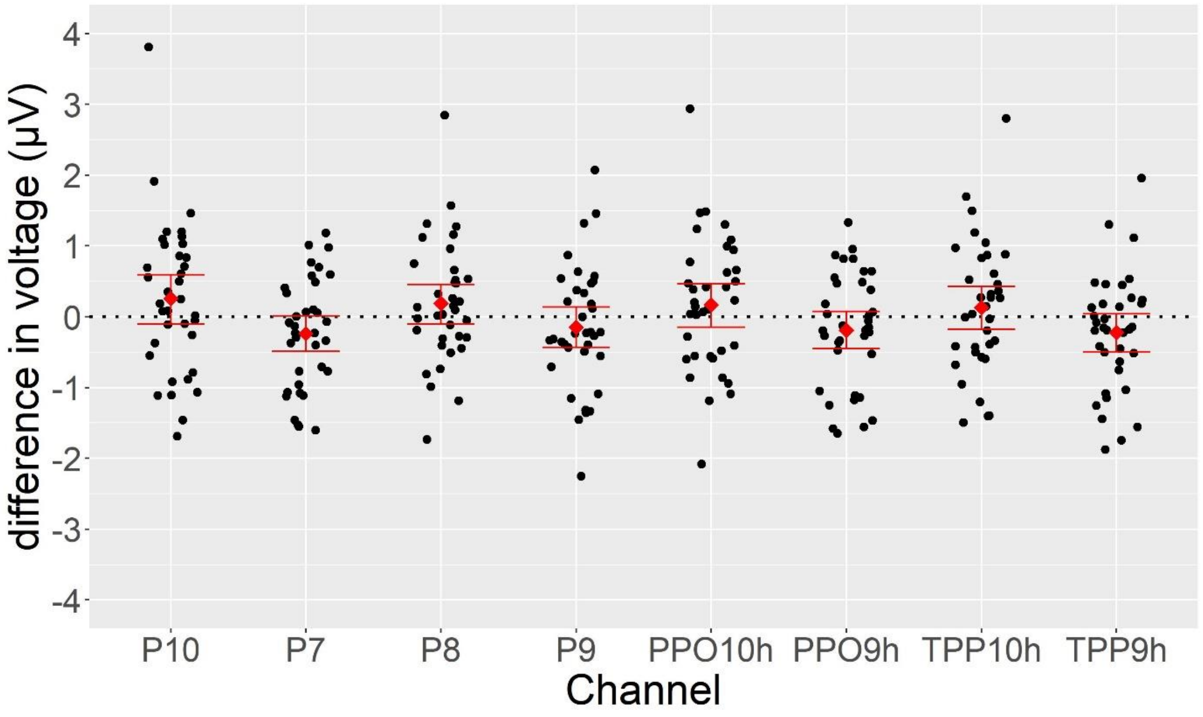


*Figure A7*. The N170 amplitude difference (texture - shape) in round-cubic difference in selected channels. Red squares represent mean estimate across participants. Error bars represent 95% percentile bootstrap confidence intervals (10000 samples) of the mean estimate.

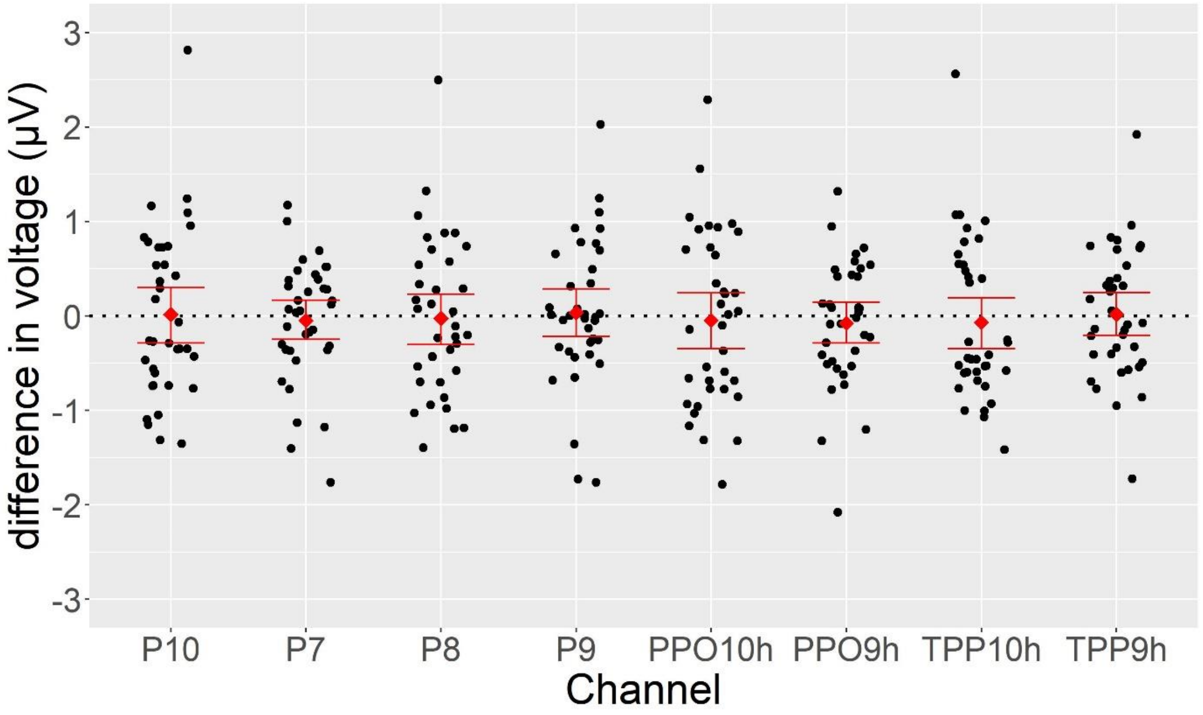


*Figure A8*. The N250 amplitude difference (texture - shape) in round-cubic difference in selected channels. Red squares represent mean estimate across participants. Error bars represent 95% percentile bootstrap confidence intervals (10000 samples) of the mean estimate.

**Plots of difference in decoding accuracy with individual data superimposed**

Figure A9 and A10 showed the same data with Figure 11 and 12, with data from individual participants superimposed.

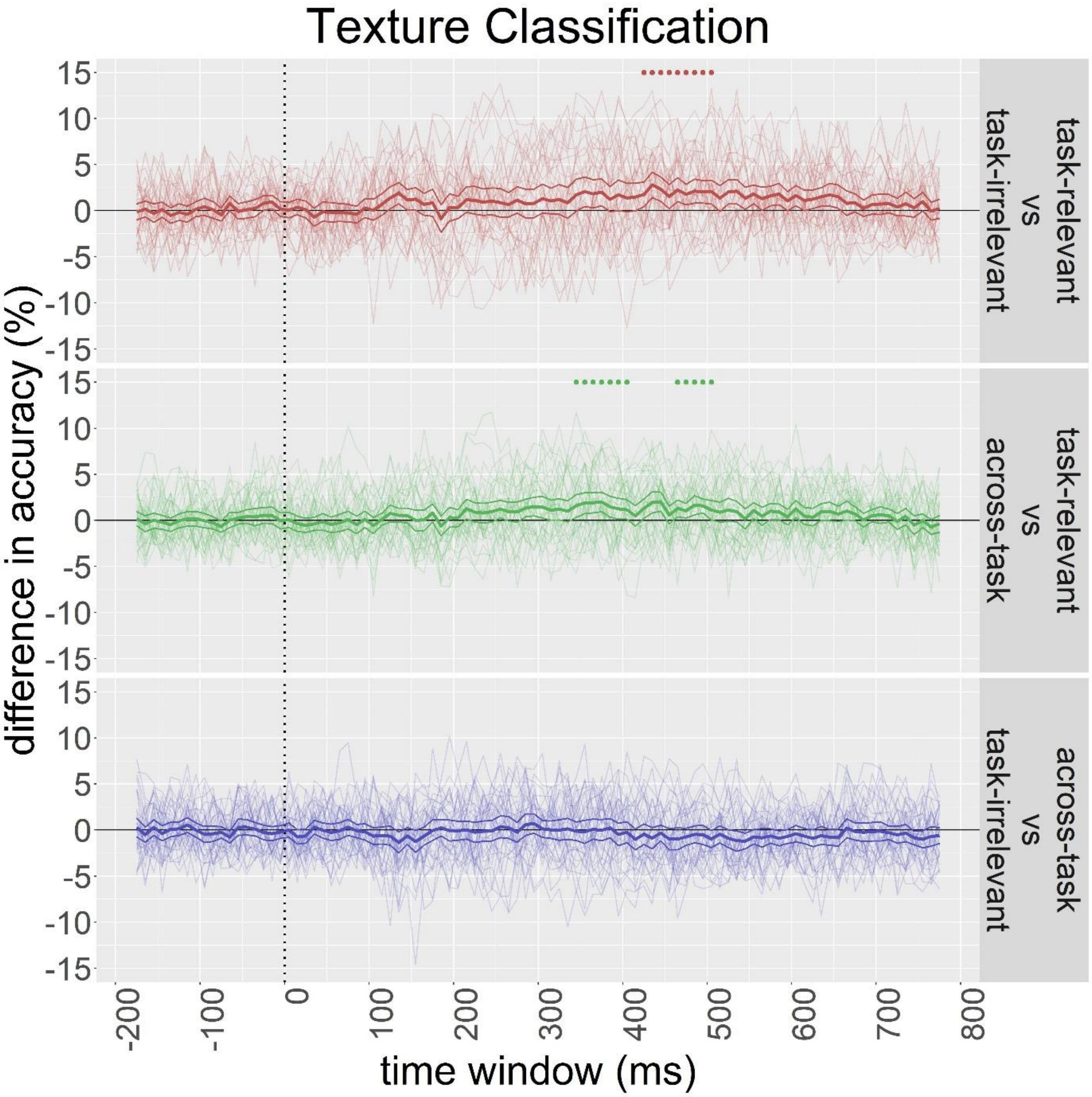


*Figure A9.* Mean difference of decoding accuracies in the comparison of task-relevant vs task-irrelevant classifiers (top panel), task-relevant vs across-task classifiers (middle panel), and task-irrelevant vs across-task classifiers (bottom panel) for texture classification with individual plots superimposed. Thinner curves around the grand average plot shows 95%

percentile bootstrap confidence intervals (10000 samples) of the mean. Dots at the top of the plots show time windows that the mean decoding accuracies of the two classifiers were significantly different.

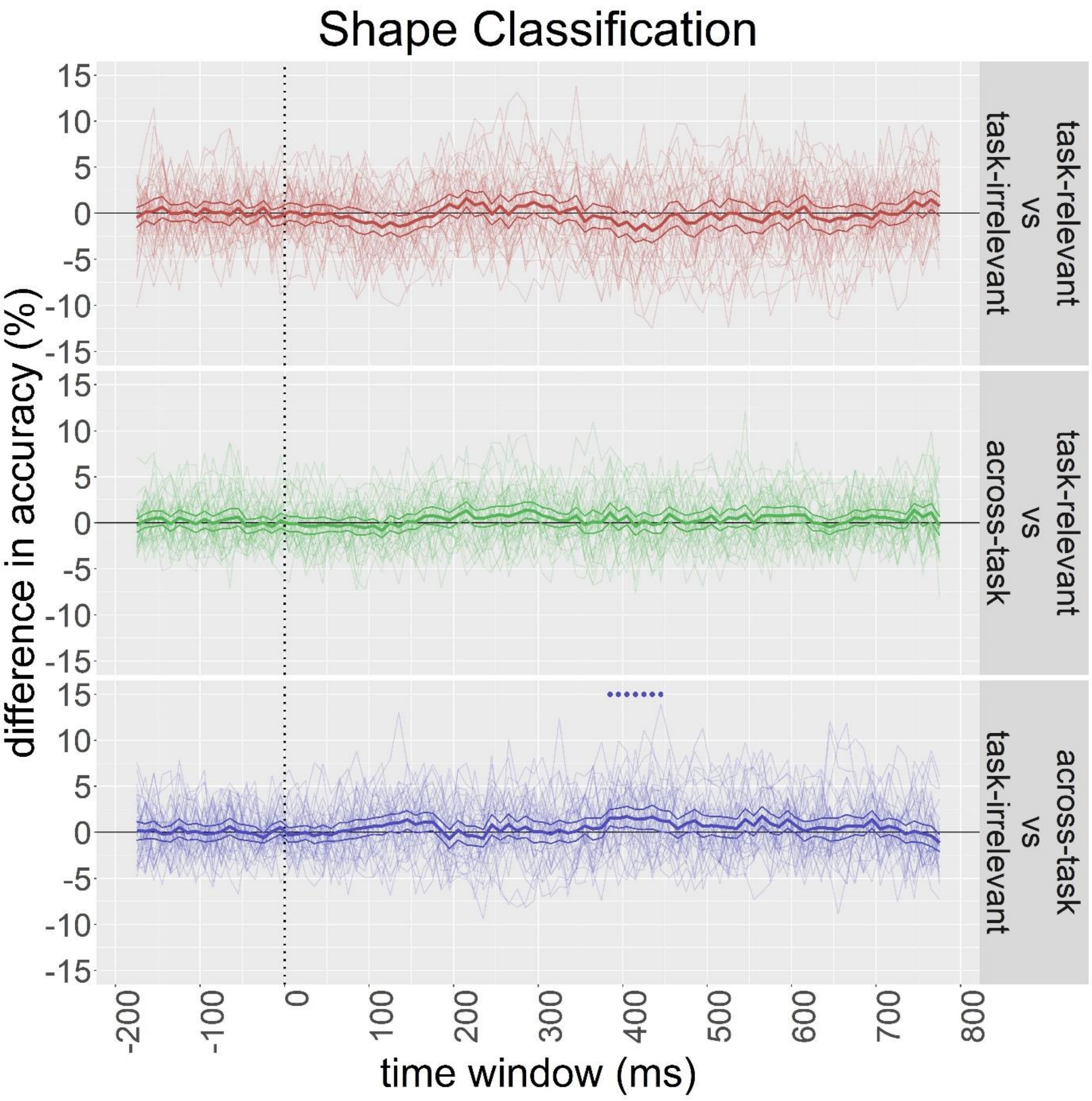


*Figure A10.* Mean difference of decoding accuracies in the comparison of task-relevant vs task-irrelevant classifiers (top panel), task-relevant vs across-task classifiers (middle panel), and task-irrelevant vs across-task classifiers (bottom panel) for shape classification with individual plots superimposed. Thinner curves around the grand average plot shows 95% percentile bootstrap confidence intervals (10000 samples) of the mean. Dots at the top of the plots show time windows that the mean decoding accuracies of the two classifiers were significantly different.

### Decoding with frontal electrodes

We conducted decoding analysis on all frontal electrodes that were excluded from the main analysis. Figure A11 and A12 show the decoding accuracies of task-relevant, task-irrelevant and across-task classifiers in texture and shape classification respectively. All classifiers in texture classification did not reach significance against chance during the whole epoch and there was no apparent peak. Decoding accuracies of classifiers in shape classification were significantly higher than chance as early as 80-130ms time window. Although peaks were not obvious in the plots, the maximum values during 100-200ms were 52.00% (task-relevant), 52.63% (task-irrelevant) and 51.77% (across-task) respectively.

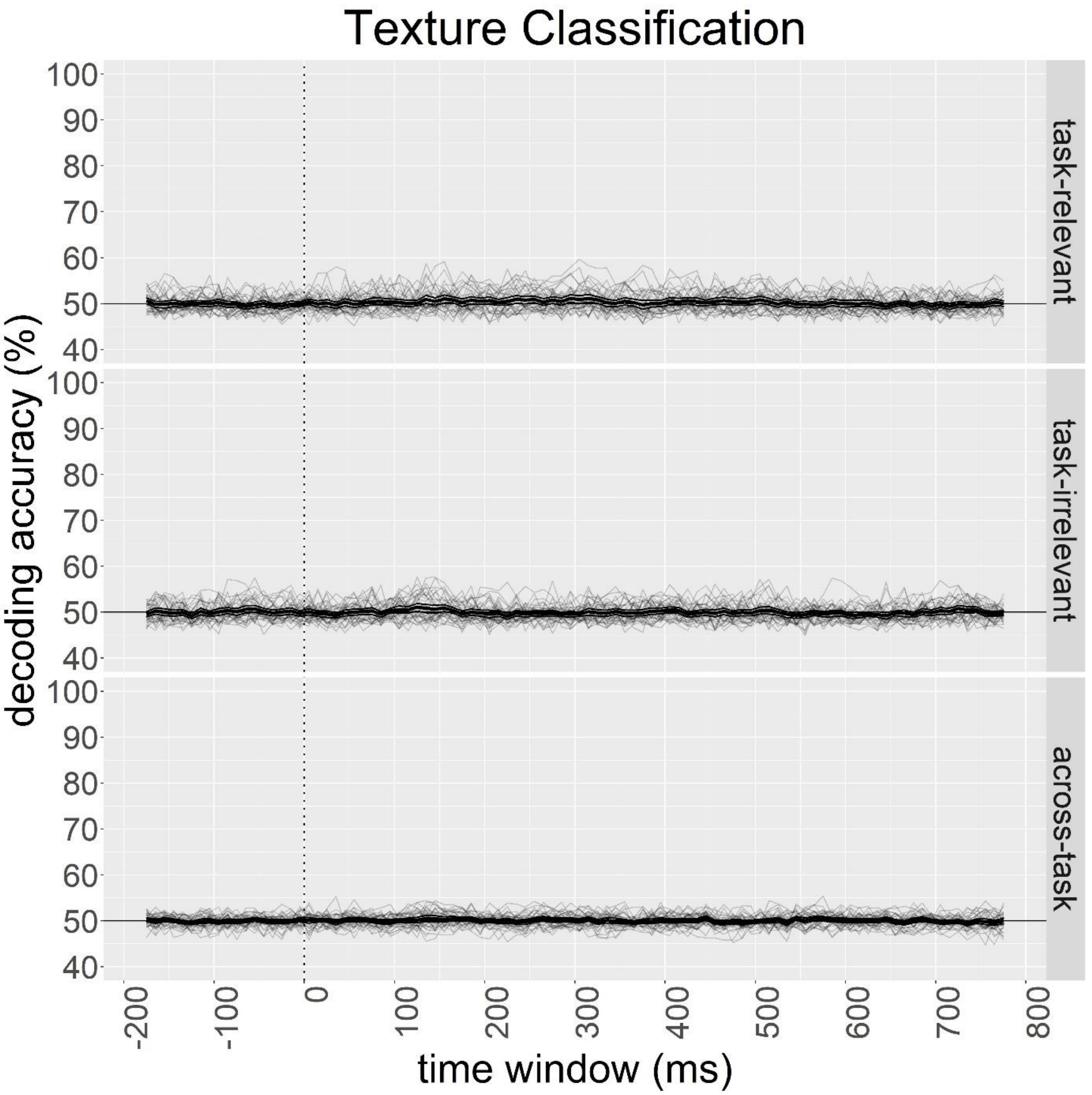


*Figure A11.* Mean decoding accuracies of task-relevant (top panel), task-irrelevant (middle panel) and across-task classifiers (bottom panel) for texture classification with individual plots superimposed. Thinner curves around the grand average plot shows 95% percentile bootstrap confidence intervals (10000 samples) of the mean. Dots at the top of the plots show time windows that the mean decoding accuracy was significantly different from chance (i.e. 50%).

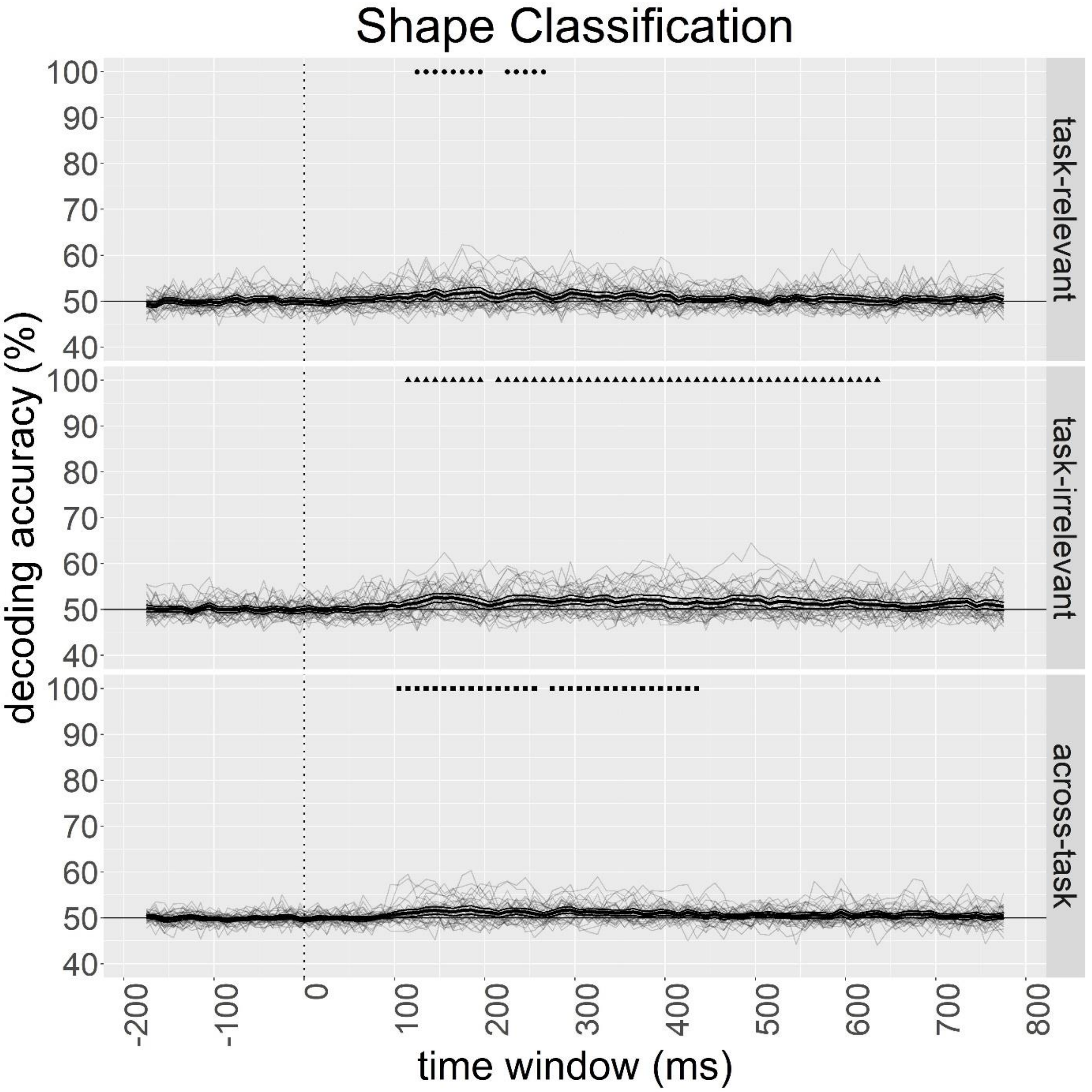


*Figure A12.* Mean decoding accuracies of task-relevant (top panel), task-irrelevant (middle panel) and across-task classifiers (bottom panel) for shape classification with individual plots superimposed. Thinner curves around the grand average plot shows 95% percentile bootstrap confidence intervals (10000 samples) of the mean. Dots at the top of the plots show time windows that the mean decoding accuracy was significantly different from chance (i.e. 50%).

Figure A13 and A14 show the comparison of classifiers in texture and shape classification respectively. The comparisons were mostly insignificant, except 200-280ms (mean = 1.22%) in task-relevant vs task-irrelevant in texture classification and 440-530ms (mean = 1.56%) in task-irrelevant vs across-task in shape classification. However, given that the decoding accuracies of individual classifiers were not significantly above chance in these periods, it is hard to interpret the results in comparisons.

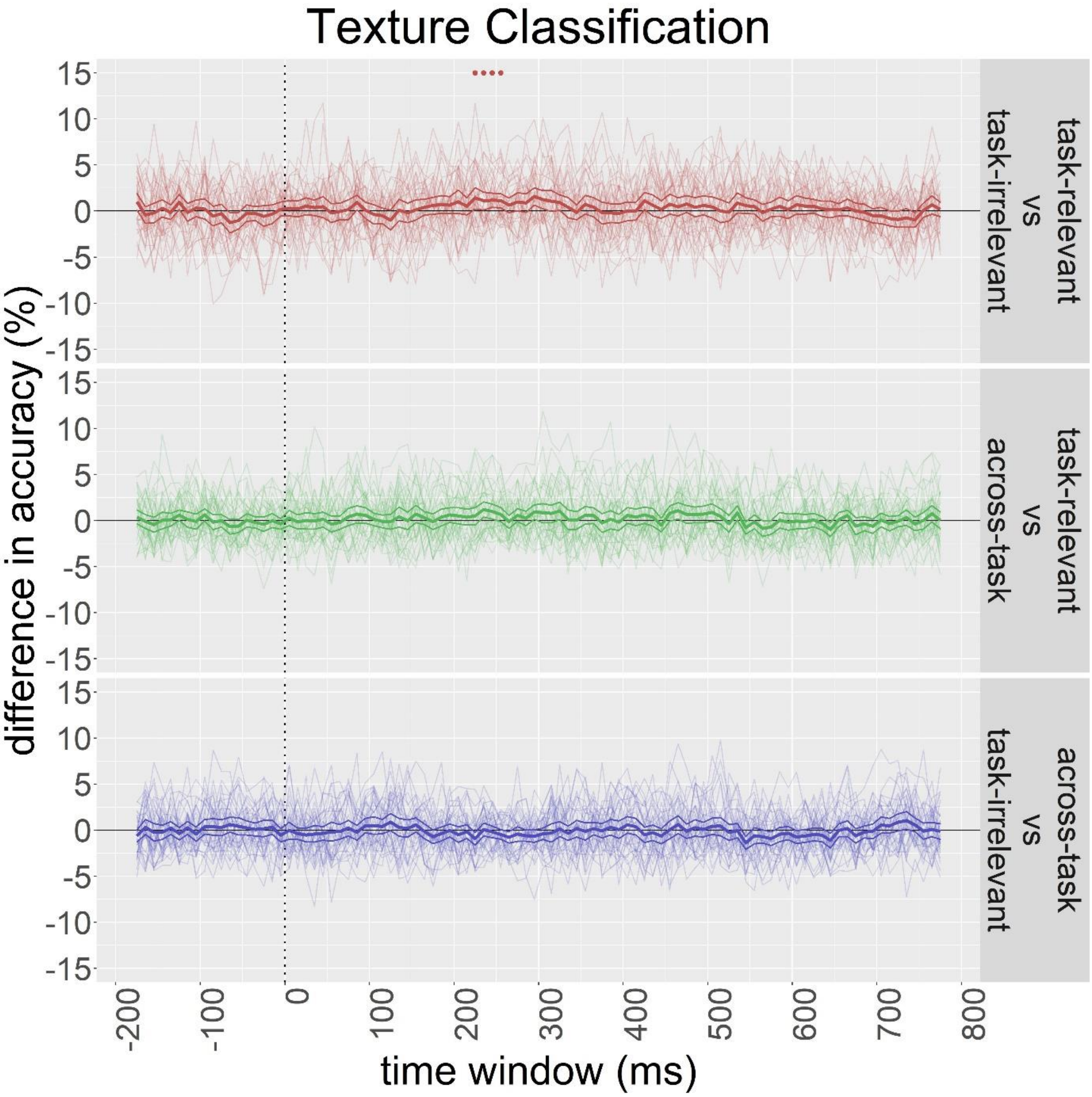


*Figure A13.* Mean difference of decoding accuracies in the comparison of task-relevant vs task-irrelevant classifiers (top panel), task-relevant vs across-task classifiers (middle panel), and task-irrelevant vs across-task classifiers (bottom panel) for texture classification with individual plots superimposed. Thinner curves around the grand average plot shows 95% percentile bootstrap confidence intervals (10000 samples) of the mean. Dots at the top of the plots show time windows that the mean decoding accuracies of the two classifiers were significantly different.

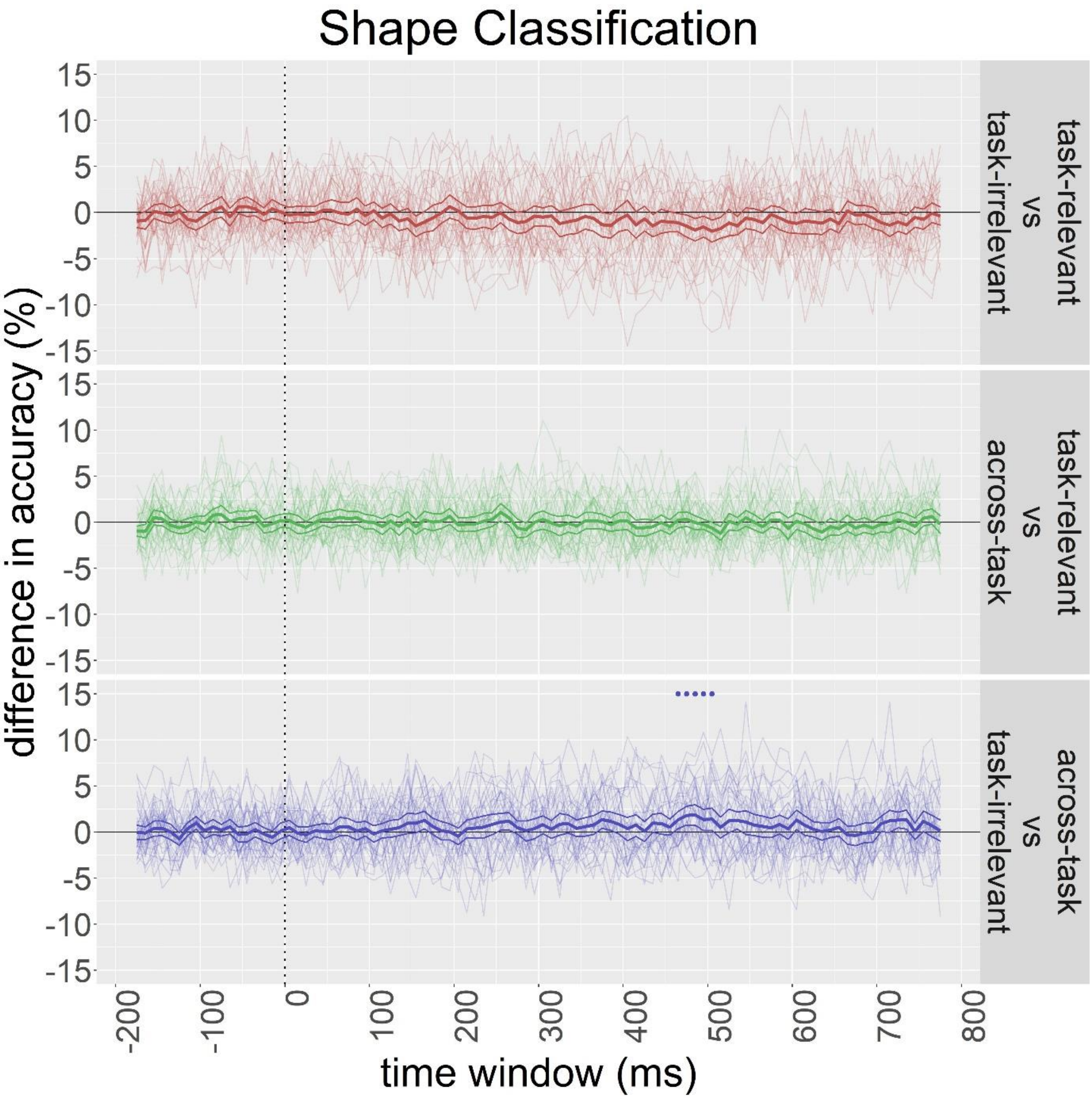


*Figure A14.* Mean difference of decoding accuracies in the comparison of task-relevant vs task-irrelevant classifiers (top panel), task-relevant vs across-task classifiers (middle panel), and task-irrelevant vs across-task classifiers (bottom panel) for shape classification with individual plots superimposed. Thinner curves around the grand average plot shows 95% percentile bootstrap confidence intervals (10000 samples) of the mean. Dots at the top of the plots show time windows that the mean decoding accuracies of the two classifiers were significantly different.
